# A distinct signature of interferon-stimulated genes linked to cross-protection against secondary viral infections in primary bronchial epithelial cells

**DOI:** 10.64898/2026.06.30.735481

**Authors:** A Tymchenko, M Fierville, P Esteves, P Gourdan, S Germain, R Ben Sghaier, M Faure, B Roger, F Rayne, N Landrein, V Magnone, P Barbry, P Berger, F Beaufils, L.E Zaragosi, H Wodrich, T Trian

## Abstract

Respiratory viral infections, such as those caused by rhinovirus, adenovirus, influenza, respiratory syncytial virus (RSV), and SARS-CoV-2, represent a major global health challenge. Despite extensive research, effective and specific antiviral treatments for these infections are still lacking, with patient care often limited to symptomatic relief. The SARS-CoV-2 pandemic, alongside the emergence of avian flu H1N1 and Nipah virus, has underscored the critical role of the respiratory tract as a critical viral target. Respiratory viral infections exhibit marked variability in infectivity and disease severity across different age groups. Epidemiological and cell-based evidence highlights distinct impacts on pediatric and adult populations. For instance, the COVID-19 disproportionately affected the elderly, while viruses like rhinovirus and adenovirus often cause severe morbidity in children. Additionally, clinical studies indicate that a primary respiratory infection can provide transient protection against subsequent infections by the same or different respiratory viruses.

In this study, we utilized a differentiated bronchial epithelial (BE) model derived from pediatric and adult donors to assess age-dependent differences under resting conditions and during viral infections. We investigated how donor age influences infection susceptibility and viral spread within the BE, focusing on the transcriptional response to rhinovirus types A and C, and adenovirus type 5. Importantly, we demonstrate that prior viral infection confers protection against subsequent infections, regardless of donor age or the initial virus type. This cross-protection is driven by interferon signaling, leading to the expression of a narrow and specific set of interferon-stimulated genes (ISGs) in both infected and bystander cells. Notably, *IFI44L* shows the strongest correlation with the level of cross protection and that its overexpression alone significantly reduces viral infection of BE. These findings suggest a distinctive, interferon-driven innate immune response profile in the BE, offering critical insights for the development of new therapeutic strategies against respiratory viral infections.

## Introduction

The respiratory tract is a major target for viral infections, which represent one of the leading causes of death globally. Although common respiratory viruses causing respiratory infections such as rhinovirus, adenovirus, flu or RSV have been well described, effective and specific treatments are currently lacking. Thus, patient care is almost always based on symptomatic treatments. The recent SARS-CoV-2 pandemic along with the emergence of viruses such as avian flu H1N1 and Nipah virus underscores the importance of the respiratory tract as key target of viral infection ^1–3^. Understanding the mechanisms of respiratory infections is essential for reducing their incidence and developing effective antiviral strategies.

A common feature of many respiratory viral infections is that infectivity and disease severity differ between age groups. Epidemiology and cellular evidence demonstrate age-dependent differences in respiratory infections, most strikingly during the Covid-19 pandemic, where the elderly experienced high mortality rates while children were largely spared ^4^. Other major respiratory viruses targeting the bronchial epithelium (BE) such as endemic seasonal rhinovirus and adenovirus, show an opposite pattern, with the pediatric population experiencing disproportionally high morbidity compared to adults^5–8^. Furthermore, several clinical studies indicate that a primary respiratory infection can transiently protect against subsequent infection by the same or other respiratory viruses^9,10^. The recent Covid-19 pandemic, characterized by a high case burden, showed that individuals previously infected with SARS-CoV-2 had a lower risk of reinfection. Similar findings were documented prior to the pandemic and subsequently reinforced by a recent systematic review ^11^. Gopal *et al.* compared the prevalence of sequential viral infections in patients with previously documented positive versus negative swabs across multiple studies. In their meta-analysis, they found a reduced prevalence of sequential viral infection following an initial positive viral swab and proposed a transient cross protective state of the respiratory tract driven by broad, non-specific innate immune mechanisms rather than virus-specific adaptive immunity. However, mechanisms regulating such a protection remain unknown and may be critical for the development of new therapeutic tools against respiratory viral infections. One of the most important family of cytokines produced during viral infection are interferons, considered as an effective, rapid and pleiotropic innate antiviral mechanism, which may contribute to the observed cross protection. Interferon signaling originates from the infected cells through pathogen sensing and propagates through autocrine and paracrine secretion to alert non-infected bystander cells and tissues^12^. Interferon signals through specific receptors and triggers the expression of Interferon stimulated genes (ISGs), which restrict viral replication and help establish a broad antiviral state in both infected and neighbouring non-infected cells^12^.

Here, we used a differentiated bronchial epithelial (BE) model derived from pediatric and adult donor samples to first assess age-dependent differences under resting physiological conditions. We then determined how donor age influences infection susceptibility and viral spread within the BE and characterized the transcriptional response following infection with rhinovirus type A, rhinovirus type C and adenovirus type 5 representing three major respiratory viruses. Finally, we explored the cross-protective effects of repeated viral infections and the underlying mechanism. Our data do not reveal age-related morphological differences in the BE under basal conditions. Instead, we show that a prior viral infection confers protection against subsequent infection by the same or a different virus, indicating an intrinsic capacity of the BE to retain a memory of infection. This protective effect is independent of donor age and of the identity of the initial virus, although we observed kinetic differences related to the spread of the primary infection. Furthermore, we demonstrate that this cross-protection is mediated by interferon induction in infected cells, leading to the expression of a select set of interferon-stimulated genes (ISGs) in both infected and bystander cells, suggesting a distinctive, interferon-driven innate immune response profile in the BE.

## Results

### Adenovirus and rhinovirus exhibit distinct infection kinetics in primary bronchial epithelium from both adult and pediatric donors

To study respiratory viral infections, we first established a primary bronchial epithelium (BE) model derived from healthy adult and pediatric donors (Table 1). To complete all experiments, basal cells were isolated from a total of 43 adults and 30 pediatric patients. Basal cells were expanded and subsequently cultured at air–liquid interface to promote differentiation into BE (Fig. 1). Full differentiation was confirmed by immunostaining for cell-specific markers of multiciliated, goblet, and basal cells (Fig. 1A), demonstrating the pseudostratified organization and presence of all three major BE cell types, with no noticeable differences between age groups. We then assessed ciliary beating (Fig. 1B) and transepithelial electrical resistance (TEER, Fig. 1C), which reflect mucociliary function and epithelial barrier integrity, respectively. All BE cultures exhibited physiological ciliary activity and robust epithelial integrity, with no significant differences between those derived from pediatric and adult donors. These findings indicate that our differentiation protocol reliably produces BE cultures with appropriate physiological properties, independent of donor age.

**Figure 1.**
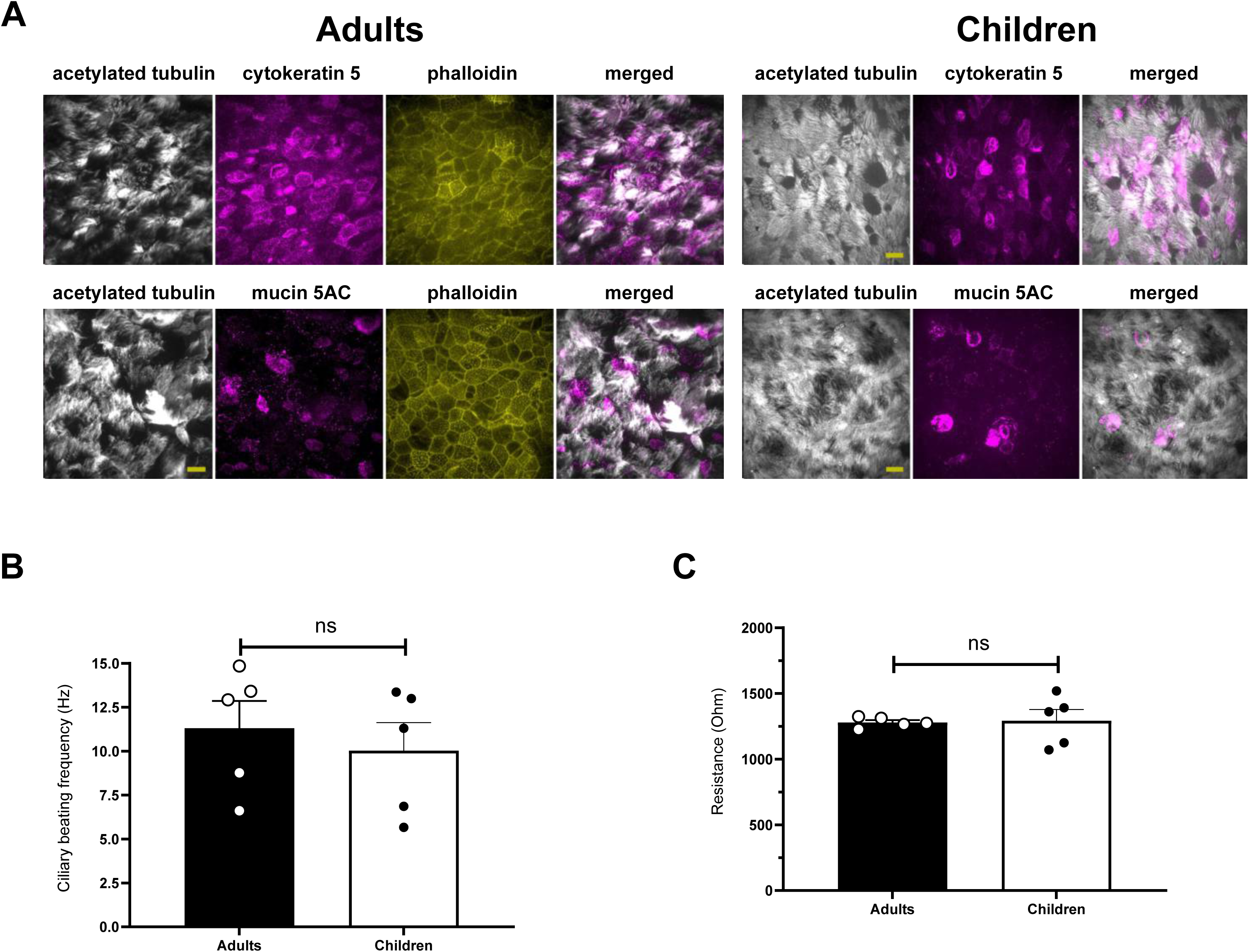
**A.** Characterization of primary BE model derived from adult (left) or pediatric (right) donors. Immunostaining of BE was performed using antibodies to cell type markers. *Acetylated tubulin* - ciliated cells marker, *cytokeratin –* basal cells marker, *mucin 5AC* – goblet cells marker. Selected area of the epithelia is shown at higher resolution (scale 10µm). **B.** Ciliary beating frequency of non-infected adult and pediatric BE. Results are presented as mean ± SEM, each dot indicates one donor. **C.** Trans epithelial resistance value of non-infected adult and pediatric donors derived BE. Results are presented as mean ± SEM, each dot indicates one donor.

**Table 1:** Patient’s characteristics.

|  | Adult<br>population | Pediatric<br>population |
| --- | --- | --- |
| Number of patients | 43 | 30 |
| Age (Mean ± SEM) | 66.9 ± 1.5 | 5.6 ± 0.8 |
| Sex (M/F) | 20/23 | 16/14 |
| BMI (Mean ± SEM) | 24.9 ± 0.6 | 17.4 ± 0.5 |
| FEV1% (Mean ± SEM) | 82.5 ± 4 | NA |

*Plus or minus values are means ± SEM. FEV1, forced expiratory volume in 1 s; BMI, body mass*
*index.*

We next investigated how differentiated BE cultures respond to infection with adenovirus (HAdV-C5, a clinical human adenovirus type 5 isolate, hereafter referred to as AdV) or rhinovirus type A (RVA) and type C (RVC). All three viruses are established respiratory pathogens with reported differences in clinical/epidemiological infection patterns between adults and pediatric ^5–8^. To assess infection dynamics, we exposed BE cultures at the apical side to each virus at an optimized multiplicity of infection to achieve robust infectivity (MOI; see Methods for details) and monitored infection kinetics (Fig. 2).

**Figure 2.**
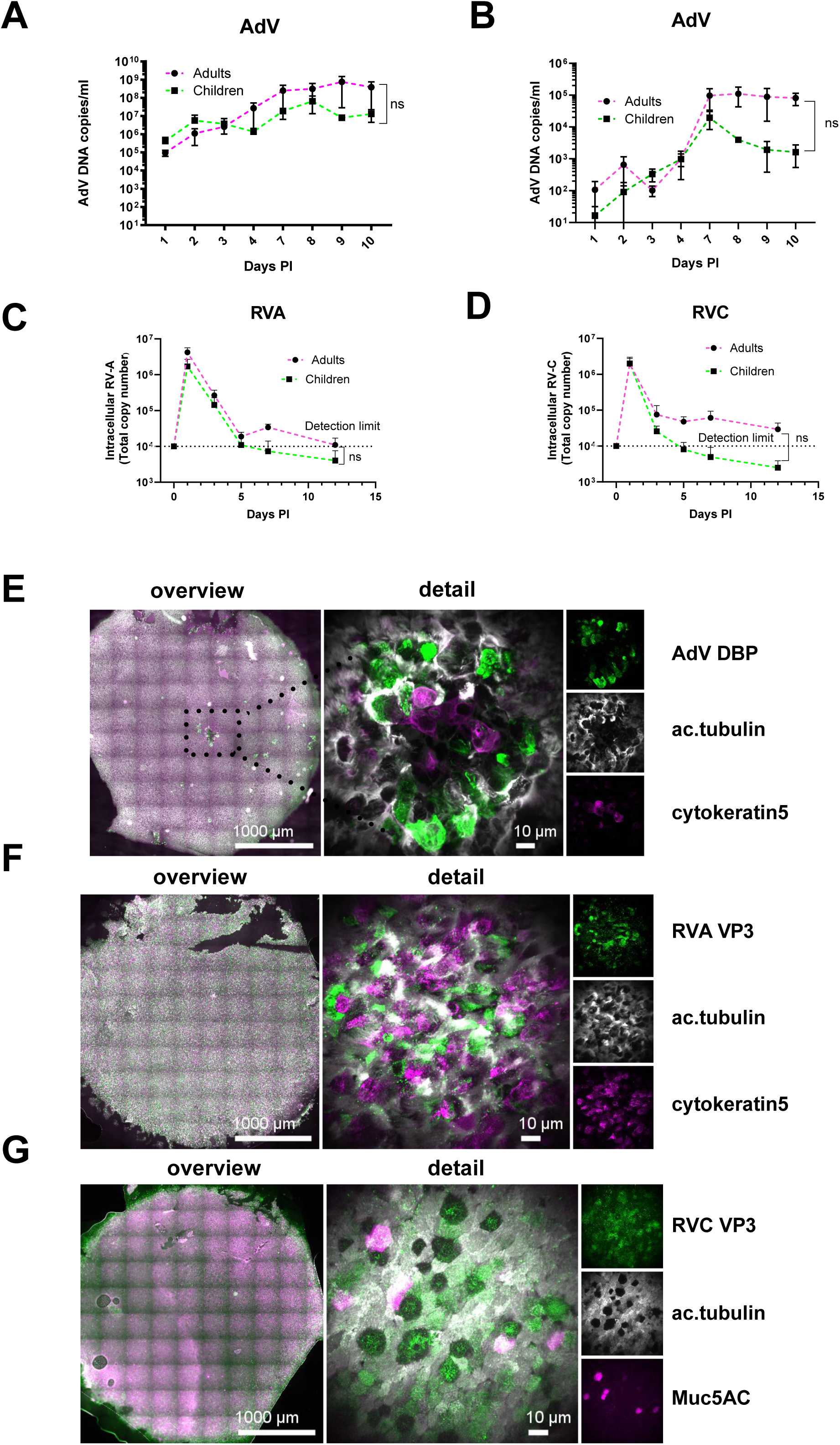
**A-B.** Ad5 infection kinetics in adult and pediatric BE. **A.** Ad5 genome release was quantified by PCR in apical side of the BE insert. **B.** Ad5 genome release quantified in basolateral side of the BE insert (n= 6 adults and 5 pediatric). **C.** RVA infection kinetics in adult and pediatric primary BE. Intracellular genome quantified was quantified by RT-PCR in whole BE (n= 7 adults and 4 pediatric). **D.** RVC infection kinetics in adult and pediatric primary BE. Intracellular genome quantified was quantified by RT-PCR in whole BE (n= 7 adults and 4 pediatric). **E.** Immunostaining of BE (adult donor) 7-days post-infection with Ad5. Immunostaining of BE was performed using antibodies to acetylated tubulin, cytokeratin and Ad5 protein DBP. Selected area of the epithelia is shown at high-resolution (scale 10µm). **F.** Immunostaining of BE (adult donor) 1-day post-infection with RVA. Immunostaining of BE was performed using antibodies to acetylated tubulin, cytokeratin and rhinovirus capsid protein VP3. Selected area of the epithelia is shown at high-resolution (scale 10µm). **G.** Immunostaining of BE (adult donor) 1-day post-infection with RVC. Immunostaining of BE was performed using antibodies to acetylated tubulin, mucin 5AC and rhinovirus capsid protein VP3. Selected area of the epithelia is shown at high-resolution (scale 10µm).

Specifically, we performed a time-course analysis, quantifying viral DNA (during 10 days for AdV) or viral RNA (during 12 days for RVA and RVC). AdV DNA quantifications in apical washes showed that AdV genomes gradually accumulated in the apical lumen (Fig. 2A), reaching a plateau by 7–10 days post infection (dpi). Around 7 dpi, AdV DNA also accumulated in the basolateral lumen, likely reflecting epithelial barrier disruption (Fig. 2B). In contrast, both RVA (Fig. 2C) and RVC (Fig. 2D) reached maximal viral loads within 24 hours of infection in the apical lumen, followed by a rapid decline to near detection limits within 3–5 dpi. RV RNA was not detected in the basolateral compartment (not shown), suggesting preserved epithelial integrity throughout infection. Despite reported age-related differences in susceptibility, we did not observe any statistically significant differences in replication kinetics for any of the three viruses between BE cultures from pediatric and adult donors. Together, the analysis shows that AdV and RV display distinct infection dynamics, with AdV reaching its infection peak, defined as maximal progeny production, around 7 dpi, whereas RV reaches maximal progeny production as early as 1 dpi.

Based on our measured kinetics, we next examined the infected BE during the respective infection peak to assess the distribution of infected versus non-infected cells (Fig. 2E-G). Adenovirus-infected cells were identified with antibodies against the DNA binding protein (DBP), which stains intranuclear replication compartments ^13^. RV-infected cells were detected using antibodies against the capsid protein VP3, staining the cytosol ^14^. To obtain a global view of infection, we performed whole-mount imaging of the entire BE and complemented this approach with higher-magnification imaging of infected regions. Co-staining of the respective viral marker with cell type–specific markers enabled identification of infected cell populations. At 7 dpi, AdV infection produced multiple, well-defined infection foci distributed across the entire epithelium (Fig. 2E). Most infected cells co-stained with CytK5 and acetylated tubulin (see inset), although occasional co-staining with MUC5AC (not shown) was observed, suggesting that AdV can infect multiple cell types. Using the same approach, we analysed RVA (Fig. 2F) and RVC (Fig. 2G) infections at 1 dpi, corresponding to the RV replication peak. In contrast to AdV, both RVs showed widespread infection across the entire BE. Co-staining with cell-type markers showed that infection was largely restricted to multiciliated cells, with rare involvement of goblet cells (see inset). All analyses were performed on at least three epithelia per age group, yielding consistent results across viruses with no apparent evidence of age-dependent differences.

### Single-Cell RNA Sequencing Uncovers Cell-Type Specific Transcriptional Responses to Adenovirus and Rhinovirus Infection in Human Bronchial Epithelium

Having characterized the replication kinetics and cell tropism of the three viruses in BE cultures, we next performed single-cell RNA sequencing (scRNA-seq) of whole infected epithelia at their respective peak infection time points (1 dpi for rhinoviruses, 7 dpi for AdV) alongside donor matched non-infected controls (Fig. 3). For each condition, defined by age group, infection status and virus type, we analyzed BE samples from three independent donors.

**Figure 3.**
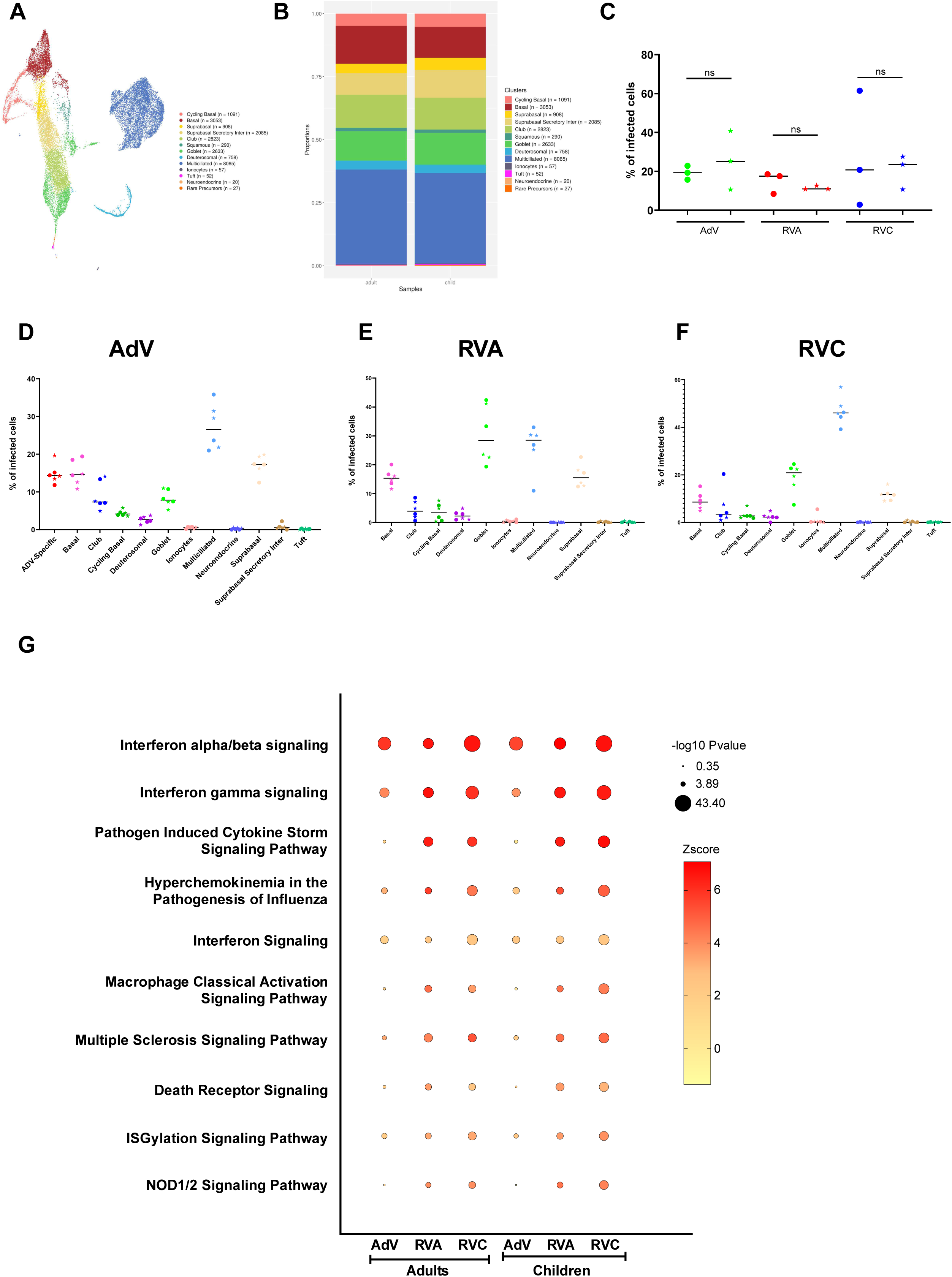
**A.** UMAP projection of the different cell types. The UMAP plots display the subset of airway epithelial cells obtained from non-infected (n= 3 adults and 3 pediatric) primary BE **B.** Proportions of each cell type in non-infected adult and pediatric primary BE. **C.** Percentage of AdV, RVA or RVC infected cells in adult (circles) and pediatric (stars) primary BE. Individual data and mean are presented. **D-F**. Distribution of infected cell in all the different cell types after AdV, RVA or RVC infection in adult (circles) and pediatric (stars) primary BE. **G.** Signaling pathways enrichment analyses were performed with human gene names using IPA platform. Comparison was performed between each BE infected condition (AdV, RVA and RVC) and non-infected BE. The size of the dots corresponds to the p-values, and color indicate the value of the z-score.

We first performed data integration of non-infected bronchial epithelial samples, to perform cell type annotation in control conditions. We identified 21,862 cells (13,256 from adult and 8,606 from pediatric donor), clustered into 13 distinct cell subtypes. In addition to the three major epithelial cell types, rare cell populations such as ionocytes and deuterosomal cells were detected (Fig. 3A and Fig. S2A) each characterized by their specific gene expression signature (Fig. S1A). The analysis confirmed full epithelial differentiation and showed a comparable cell type composition, both qualitatively and quantitatively, between the two age groups (Fig. 3B). No significant difference was detected when comparing cell type proportions between adult and pediatric donor (Fig. 3B). For each cell population, we next conducted a pseudobulk differential gene expression analysis between adult and pediatric BE, followed by gene set and pathway enrichment analysis (Ingenuity Pathway Analysis ^15^). We found 119 genes that were significantly upregulated in adult BE, compared with 42 in pediatric BE (Supplementary Table S1). However, the IPA showed that no major pathways were uniquely enriched in either age group. Taken together, these results demonstrate that BE derived from adult and pediatric donors display similar cellular composition and steady-state transcriptional activity, consistent with our morphological characterization.

We next integrated viral-infected samples together with control samples in order to analyse infection-specific effects. Umap and cell annotation of infected conditions are presented in Supplementary Figure 2. Using virus-specific transcripts (AdV, RVA, RVC), we were able to identify cells expressing viral reads, which were subsequently used to determine the infected cell population based on pre-defined thresholds (see materiel and methods section). We first compared the percentage of infected cells between adults and pediatric BE for each virus, merging all cell types (Fig. 3C). We did not find any significant differences between the two age groups, suggesting no difference in overall infection susceptibility which is consistent with our morphological analysis and virus release assays. We next determined which cell types were infected for each condition. In AdV-infected BE, infected cells were mainly multiciliated, basal and suprabasal cells (Fig. 3D). Moreover, we found in AdV-infected BE a novel distinct cell population characterized by a unique gene signature, which we termed *AdV-specific* cells (Fig. S2B). Interestingly, while these cells do not represent the majority of infected cells, their level of infection was >90%, a percentage much higher than what was found for multiciliated, basal and suprabasal cells, for which only about 20% of cells were infected (Fig. S3A-C). This observed heterogeneity in infected target cells is consistent with the variability we observed in our morphological analysis (Fig. 2). For RVA (Fig. 3E) and RVC (Fig. 3F) transcripts were mainly detected in multiciliated cells, which is reflecting our immunofluorescence analysis. However, we also found a subset of infected goblet, basal and suprabasal cells. Notably, the relative distribution of infected cells for all three viruses in each case were identical between pediatric and adult donors.

Viral infection also profoundly changed the transcriptional profile of infected BE compared with donor matched non-infected controls. Infection with either AdV, RVA or RVC induces a major transcriptional reprogramming in all cell types dominated by ISG expression (Fig. S4). Gene Set Enrichment Analysis using IPA confirmed substantial gene expression changes by viral infection. All three viruses elicited a robust interferon response in both adult and pediatric BE resulting in significant z-scores of IPA pathways related to interferon signaling and activation (Fig. 3G). We thus focused our analysis on the interferon response.

To determine whether the interferon response originated from specific cell types, we repeated the analysis focusing on individual cell types rather than the entire BE. For this purpose, we compiled a comprehensive list of ISGs from the Interferome database (https://interferome.org/interferome/home.jspx ^16^) and used it to identify significantly upregulated ISGs within each defined cell type (Fig. 4). First, when combining adults and pediatric samples, we observed a robust upregulation of ISGs across nearly all BE cell types, irrespective of the infecting virus, indicating that ISG induction occurred both in directly infected cells and in non-infected bystander cells (Fig. 4A). Interestingly the most robust induction was limited to a well-defined subset of ISGs (See ISGs name highlighted in Fig. 4A). This observation remained consistent when analyzing adult or pediatric BEs separately (Fig. S5A-B). Together, these analyses indicate that the BE mount a coordinated and well-defined response to viral infection that is largely independent of cell type or infection status. Given that ISGs are induced by interferons, we next asked which cells within the infected BE were responsible for interferon expression.

**Figure 4.**
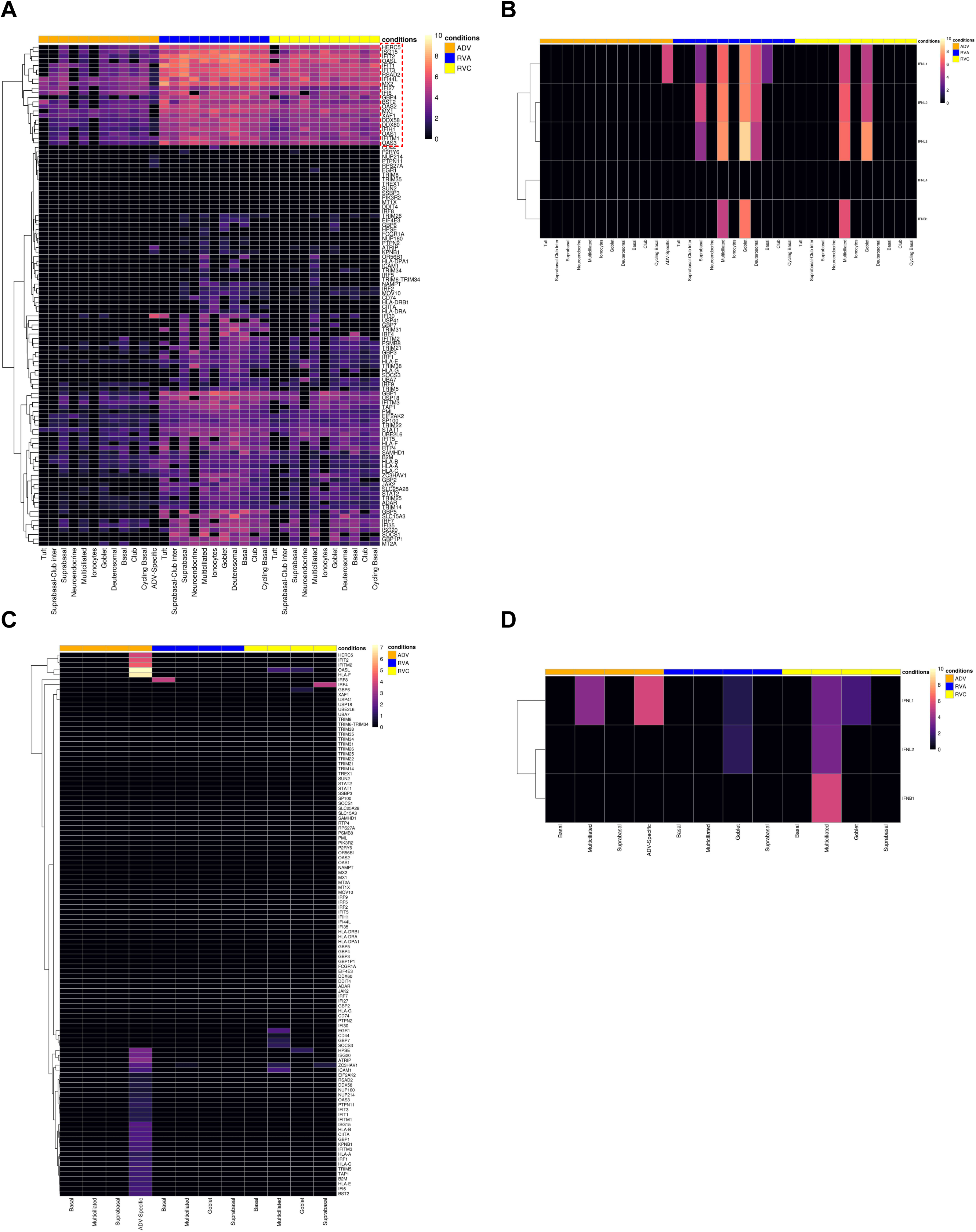
**A.** Heatmap of clustered Interferon induced genes upregulation for each cell type in merged adult and pediatric BE (n= 3 adults and 3 pediatric). ISGs activation was assessed between AdV, RVA or RVC infected primary BE compared to non-infected BE. Red frame highlight subset of induced and common ISGs **B.** Heatmap of clustered Interferon genes upregulation for each cell type in merged adult and pediatric BE. Interferon genes activation was assessed between AdV, RVA or RVC infected primary BE compared to non-infected BE (n= 3 adults and 3 pediatric). **C.** Heatmap of clustered Interferon induced genes upregulation in cell type targeted by either AdV, RVA or RVC in merged adult and pediatric BE. ISGs activation was assessed between AdV, RVA or RVC infected cells and bystander non-infected cells from the same BE (n= 3 adults and 3 pediatric). **D.** Heatmap of clustered Interferon genes upregulation in cell type targeted by either AdV, RVA or RVC in merged adult and pediatric BE. Interferon genes activation was assessed between AdV, RVA or RVC infected cells and bystander non-infected cells from the same BE (n= 3 adults and 3 pediatric).

Unlike the broadly distributed ISG expression, upregulation of interferon genes (IFNλ and IFNβ) was confined to the infected cell population, with a virus-specific pattern for each of the three infections within a given BE (Fig. 4B). We observed the same restriction of interferon expression to infected cells when analyzing adult or pediatric samples separately (Fig. S5C-D). A plausible explanation is that interferon production is initiated in infected cells, thereby eliciting a robust antiviral response in bystander cells throughout the BE. Here, bystander cells are defined as those lacking detectable viral reads despite originating from an infected BE.

To confirm this, we performed a second analysis in which infected cells and bystander cells from infected BE were compared to their respective counterpart in non-infected BE. We therefore compared ISG and interferon gene expression between infected cells and bystander cells from the same cell subtype across the three infected conditions. This analysis was focussed on the main infected subtypes for each virus identified in Figure 3. Combined analysis of adult and pediatric samples did not reveal any significant differences in ISG expression between infected and bystander cells for any of the three viruses (Fig. 4C). In this context, AdV-specific cells were the exception, as their high infection rate (>90%) precluded the identification of suitable non-infected bystander counterparts. In contrast to ISGs, interferon gene expression was markedly upregulated in infected cells compared to bystander cell of the same subtype (Fig. 4D). Similar results were obtained when we specifically performed the analysis for either adult (Fig. S5 E,G) or pediatric BE (Fig. S5F,H). This analysis confirmed robust ISG upregulation across nearly all BE cell types in both age groups, regardless of the infecting virus. In contrast, interferon gene expression was specifically upregulated in infected cells, whereas non-infected bystander cells of the same cell type showed no detectable interferon expression. Thus, interferon transcripts are enriched in virus-positive cells, likely driving paracrine interferon signaling that broadly induces ISGs and promotes an antiviral state throughout the epithelium.

#### Protection from subsequent rhinovirus infection in pre-infected epithelia occurs independently of age and viral strain, but varies with infection timing

Our analysis revealed a strong induction of a restricted subset of ISGs (following infection with each of the three viruses. Moreover, these ISGs subset partially overlapped between viruses, suggesting a common BE-specific response. ISGs are well known as potent inhibitors of viral infection that can limit both viral infection and replication. Given the overlap in ISGs we thus hypothesized that primary infection with respiratory viruses may establish a cross-protective, ISG-driven infection-refractory state that shields the epithelium from subsequent infections in a more general way. RVA was recently identified in a cross-sectional study to be the most commonly found secondary infecting virus ^11^. To test this hypothesis, we established an experimental setup in which epithelia were first infected with either AdV, RVA, or RVC, and subsequently challenged on consecutive days with a GFP-expressing RVA, enabling quantification of residual infectivity 24 h later across the entire BE (Fig. 5A).

**Figure 5.**
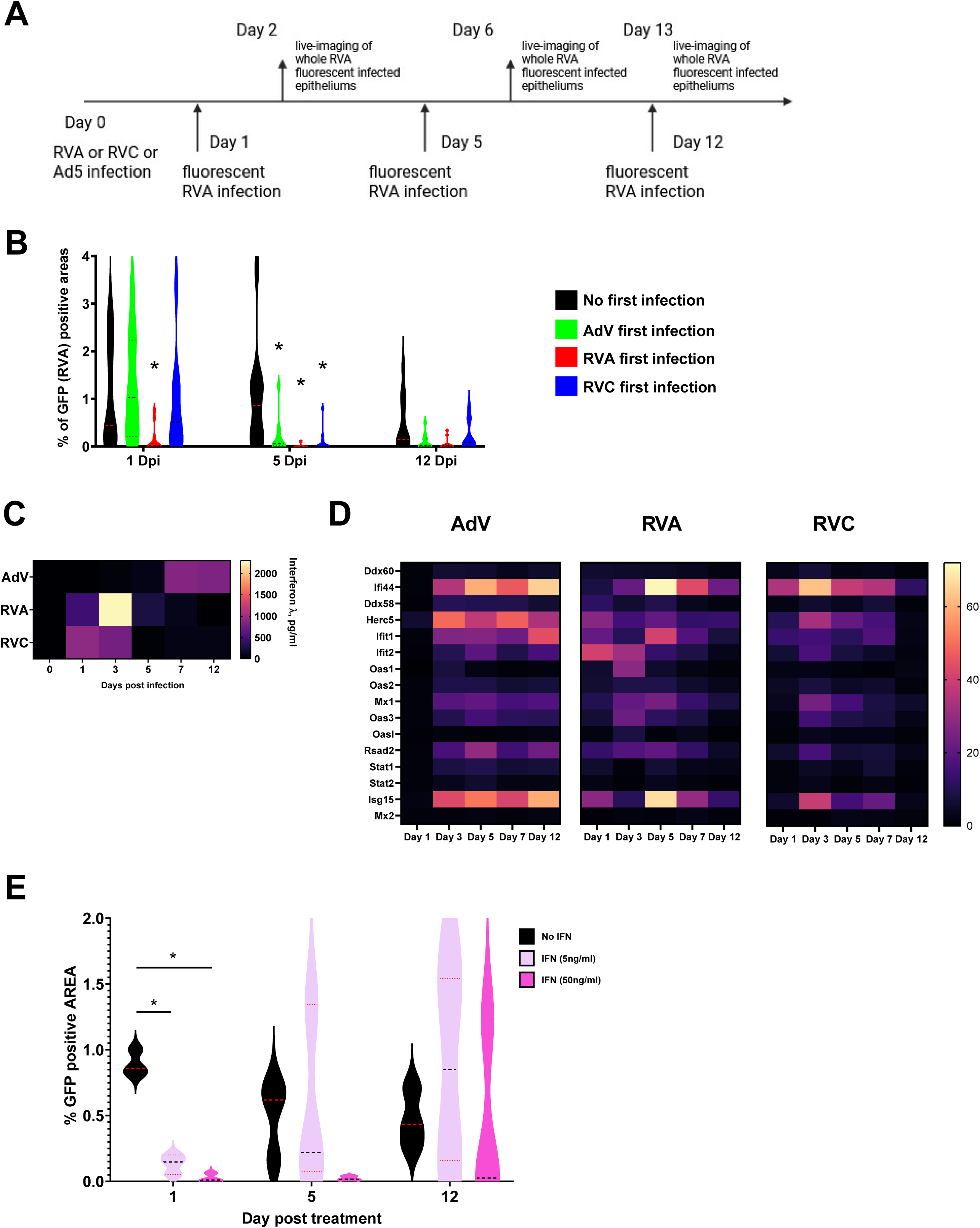
**A.** Experimental set-up design for cross-protection experiment. **B.** Percentage of GFP positive area induced by RVA-GFP infection in AdV (green), RVA (red), RVC (blue) and non-infected control (black) pre-infected adult primary BE after 1-, 5- and 12-day post infection (n= 11 adults). **C.** Heatmap of the kinetics of interferon λ release in AdV, RVA or RVC infected adult primary BE assessed by ELISA (n= 8 adults). **D.** Heatmap of the kinetics of ISGs fold increase mRNA expression in AdV, RVA and RVC infected adult primary BE assess by real time PCR (n= 7 adults). **E.** Percentage of GFP positive area induced by RVA-GFP infection in adult primary BE either after no treatment (Black) or with 5ng/ml (purple) or 50 ng/ml of interferon λ after 1, 5- and 12-day post infection (n= 4 adults).

Epithelia pre-infected with RVA, RVC, or AdV from both adult and pediatric donors exhibited varying degrees of cross-protection against subsequent RVA infection (Figs. 5B and S6A). The kinetics of this protection differed between viruses (Fig. 5B). Rhinovirus pre-infected epithelia from both age groups showed rapid cross-protection, detectable within 24 h after primary infection, peaking around day 5, and declining by day 12, the latest time point analyzed. This effect was more pronounced for RVA, with significant protection observed as early as 1 dpi. RVC followed a similar rapid cross-protection kinetics but did not reach significance in early time points: at 1 dpi, only six of eleven BEs displayed cross-protection whereas the remaining samples displayed delayed protection. AdV pre-infected epithelia showed no early cross-protection at 1 dpi. RVA infection was even slightly, though not significantly, enhanced 24 h after AdV infection. Robust AdV-mediated cross-protection emerged by day 5 and declined again by day 12, mirroring the kinetics observed with rhinoviruses. The same pattern of cross-protection was observed in pediatric BEs (Fig. S6A). Together, these experiments demonstrate that primary respiratory virus infections establish a transient cross-protective state in the bronchial epithelium that is effective against secondary infections with RVA.

We next investigated the mechanisms by which primary infections with three distinct viruses elicited a robust, yet differently timed, epithelial cross-protection. Our scRNA-seq analysis at peak infection times (1 dpi for RV and 7 dpi for AdV) revealed strong, epithelium-wide induction of a restricted and overlapping set of ISGs. Notably, ISG upregulation occurred in both infected and bystander cells, whereas IFN expression was restricted to infected cells. Based on the infection kinetics, we hypothesized that differences in IFN production and subsequent IFN responses might determine the timing of cross-protection. In this scenario, viral replication dynamics would primarily govern the induction of interferon and subsequent ISG.

To test this, we assessed the kinetics of IFN and ISG expression in response to all three viruses for both age groups within the BE. We quantified type I interferon (IFN-β) and type III interferon (IFN-λ) levels in basal medium up to 12 dpi using ELISA. While IFN-β remained undetectable in most donors (not shown), IFN-λ was readily measurable for both adult and pediatric BE (Figs. 5C and S6B). IFN-λ levels rose early for RVA- and RVC-infected epithelia, peaking at 2–3 dpi before declining. In contrast, AdV-infected epithelia displayed a delayed IFN-λ response, with levels rising from 5 dpi and reaching their peak at 7–12 dpi. Thus, IFN production in the medium closely followed the infection kinetics (Fig. 1) but declined more rapidly than the observed timeline of cross protection.

We next monitored the temporal expression kinetics of a selected panel of ISGs that were upregulated in response to each of the three viral infections in the scRNA-seq experiment (Fig. 4A). Total RNA was extracted on consecutive days following infection, and ISG expression was quantified by RT-qPCR. The temporal expression patterns of the analyzed ISGs closely paralleled those of interferon. In RV-infected epithelia, ISG expression was detectable from 1 dpi, peaked at 3–5 dpi, and gradually declined thereafter. In contrast, AdV-infected epithelia exhibited delayed ISG induction, starting at 3 dpi and progressively increasing, with robust activation still evident for most ISGs at 12 dpi (Figs. 5D and S6C). The expression kinetics were consistent across both age groups, and ISG expression followed the virus-specific replication dynamics, although absolute expression levels differed. Together, these findings pointed to a strong role of type III interferon in shaping the epithelial response. To test whether IFN alone was sufficient to confer cross-protection, we replaced the primary viral infection in our cross-protection assay with exogenous IFN treatment (Fig. 5E). Epithelia from the same donors were pre-treated with two concentrations of IFN-λ (5 or 50 ng/ml per epithelium) and subsequently challenged with RVA-GFP at 1-, 5-, or 12-days post-treatment. Cross-protection was assessed as before (Fig. 5A). We observed rapid, dose-dependent protection as early as 24 h post-treatment, confirming that IFN can induce a fast and protective state. However, this effect declined by day 5, with only residual protection remaining at the higher IFN dose. Notably, the high IFN dose was ∼10-fold higher than levels measured during viral infection, yet conferred protection comparable to, or weaker than, that induced by infection itself. Taken together, these findings suggest that efficient cross-protection requires either sustained interferon signaling or the contribution of additional pathways. Both mechanisms are likely to be more effectively engaged during ongoing viral infection than by a single application of IFN.

To identify molecular factors that might underlie the cross-protection mechanism, we integrated matched cross-protection data (Fig. 5B) with corresponding RNA sequencing profiles using BE from five individual donors (Fig. 6). To minimize donor-to-donor variability in infection levels and cross-protection, we quantified, for each donor, cross-protection as the GFP-positive surface area normalized to that of non-infected BE from the same donor at the corresponding time point (normalized data are shown in Fig. S7A). For each cross-protection time point, we collected RNA from the corresponding BE cultures and performed bulk RNA sequencing. This approach enabled parallel analysis of cross-protection and associated transcriptional profiles at the individual donor level.

**Figure 6.**
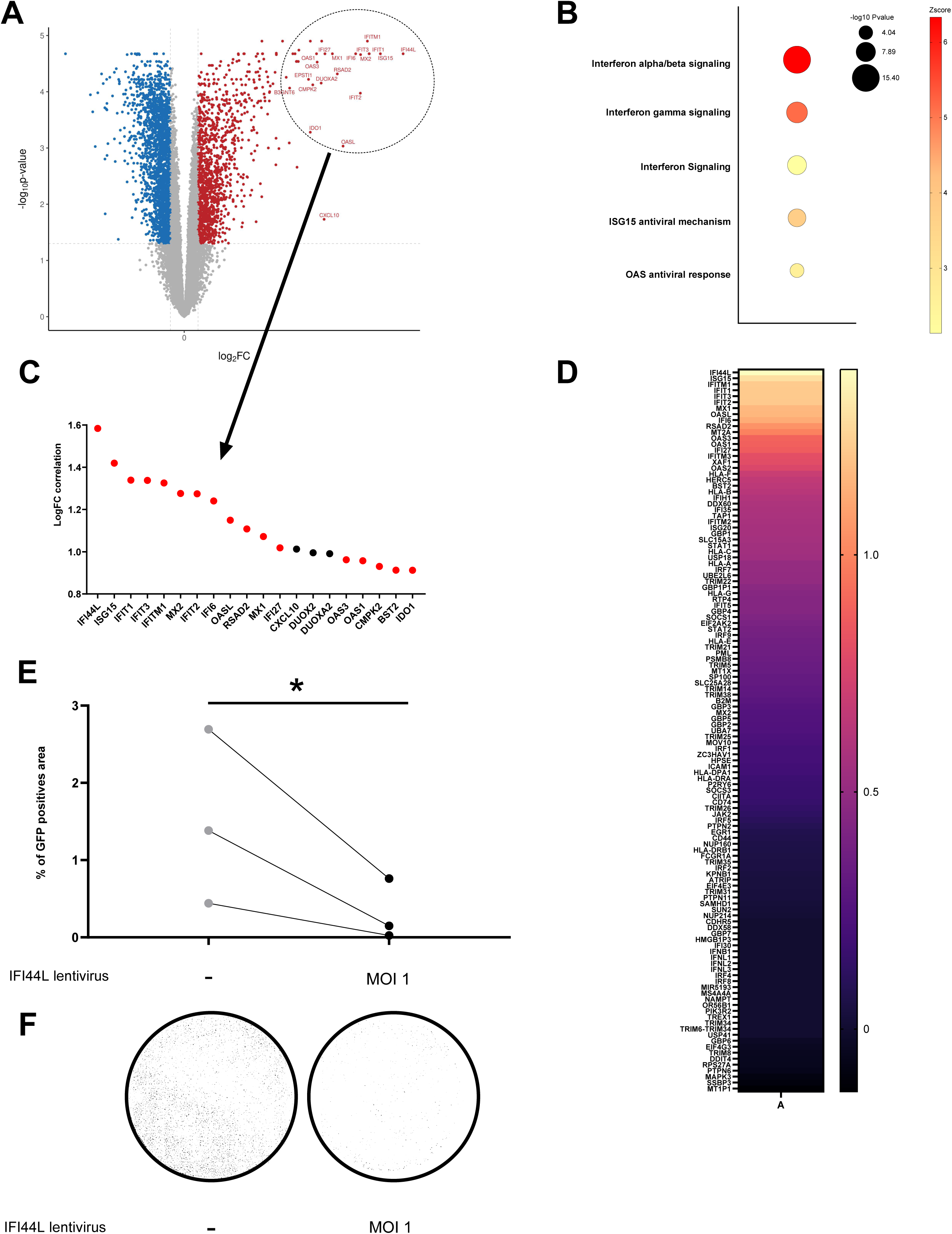
**A.** Volcano plot of genes identified with bulk RNA sequencing associated to corresponding cross protection level analyzed with limma linear regression model. Red dots are genes significantly positively correlated with cross-protection and blue dots are genes significantly negatively correlated with the cross-protection. Non-significant genes are depicted as grey dots. **B.** Signaling pathways enrichment analyses were performed with human gene names using IPA platform on genes significantly positively correlated (red dots from Figure 6A). The size of the dots corresponds to the p-values, and color indicate the value of the z-score. **C.** Top 20 genes positively correlated with cross-protection from the limma linear regression model. Red dots indicate ISGs. **D.** Heatmap of correlation value for all ISGs identified with bulk RNA sequencing. **E.** Percentage of GFP positive area induced by RVA-GFP infection in non-infected control (grey dots) or in IFI44L expressing lentivirus (black dots) BE. **F.** Representative images of GFP signal induced by RVA-GFP infection (white) in non-infected control or in IFI44L expressing lentivirus BE.

The RNA sequencing data were first used to assess infection kinetics by viral reads quantification. Consistent with our previous results (Fig. 2), AdV infection showed a low infection rate at day 1, reaching a peak at day 5 that remained detectable at day 12. In contrast, RVA and RVC showed early infection peaks at day 1 followed by a rapid decline towards the detection level at day 5, with no detectable infection at day 12 (Fig. S7B). Moreover, using cell deconvolution of the bulk RNA-seq data informed by the scRNA-seq data, we assessed the proportion of cell types within each sample to exclude infection-induced major changes in BE composition over time (Fig. S7C). Overall, epithelial cells composition remain largely unchanged after infection, with only a modest reduction in multiciliated cells in RVA-infected cultures at 5dpi.

We then analyzed the significantly and differentially expressed genes in each cross-protection condition compared to the associated non-infected controls. We observed gene expression changes induced by all viruses at every time points except for AdV at 1 dpi. Interestingly, the twenty most overexpressed genes in each condition were almost exclusively ISGs (Fig. S8A). Consistent with this, IPA analysis identified interferon pathway activation among the five strongest induced pathways (Fig. S8B). Moreover, the temporal pattern of ISG (Fig. S9A) and interferon (Fig. S9B) expression displayed virus specific kinetics. In RV-infected epithelia, ISG expression increased at 1 and 5 dpi before declining at day 12. In contrast, AdV-infected epithelia showed delayed ISG induction, beginning at 5 dpi which continued to rise with robust activation still evident for most ISGs at 12 dpi. Taken together, the bulk RNA sequencing confirmed our finding of a cross BE interferon response controlled by virus propagation kinetics.

To identify common determinants of cross-protection across the three viruses, we next applied limma, a linear regression model, to integrate the RNA sequencing data with the cross-protection levels and thereby to identify genes which would be significantly correlated with cross-protection ^17^. This analysis revealed 1324 genes, which were significantly positively correlated and 2185 genes that were significantly negatively correlated with cross protection (Fig. 6A). Ingenuity pathway analysis revealed that most genes positively correlated with cross-protection were also strongly associated with the interferon response, as reflected by highly significant z-scores for interferon signaling and activation pathways (Fig. 6B). This finding was further supported by the observation that, among the twenty genes showing the strongest positive correlation, fourteen were ISGs, including nine out of the top ten (Fig. 6C). Importantly, not all ISGs showed a strong positive correlation. When we restricted our analysis to upregulated ISGs detected in the RNA sequencing data, we found that the correlation between general ISG expression and cross-protection was confined to a narrow subset, revealing a distinct and highly specific ISG signature, with IFI44L showing the strongest correlation with cross-protection (Fig. 6D). To validate a potential role for IFI44L in BE cross protection, we generated a lentiviral vector expressing IFI44L together with “mCherry” separated by an IRES element and transduced basal epithelial cells during the initial seeding phase on cell culture inserts. Transduced cells differentiated into BE with normal kinetics and morphology. Following complete differentiation, we observed within the BE multiple transduced cells identified through expression of the fluorescent marker (Fig. S9C) and confirmed ectopic expression of IFI44L in transduced BE using western blotting although expression levels were low (Fig. S9D). To measure a potential effect of IFI44L overexpression we repeated the cross-protection experiment comparing RVA-GFP infection in IFI44L transduced BE vs. un-transduced BE. Our analysis revealed that IFI44L overexpression consistently and significantly reduced RVA infection (Fig. 6E-F). Moreover, none of the infected cells (marked by GFP) colocalized with “mCherry” positive cells (data not shown) suggesting a possible antiviral effect mediated through IFI44L at the cellular level.

Together, our results identify a selective interferon-stimulated gene signature that is strongly associated with bronchial epithelial cross-protection. They further indicate that antiviral resistance in the airway epithelium is driven by a defined subset of ISGs, rather than a broad, global interferon response. Notably, this conserved ISG signature is induced by viruses of diverse origins with distinct replication strategies and infection kinetics, underscoring a shared epithelial antiviral defense mechanism.

## Discussion

Clinical observations and epidemiological studies clearly indicate that children and adults respond differently to common respiratory viruses. As an example, the Covid-19 pandemic has revealed much higher morbidity in the elderly population following a SARS-CoV-2 infection compared to children. For common cold causing viruses such as adenovirus and rhinovirus the opposite pattern is observed, with a substantially higher disease burden in the pediatric population ^6,8^. Moreover, within the rhinovirus family, RV-C shows a particularly strong association with pediatric asthma exacerbations. These observations raise the possibility that pediatric and adult airway epithelia differ in their intrinsic susceptibility or antiviral responses. To address this question, we compared infections by adenovirus (AdV), RV-A, and RV-C in differentiated human bronchial epithelia derived from pediatric and adult donors. Despite their similar epidemiological profiles in children, AdV and RV represent fundamentally different infection strategies and infection kinetics. AdV establishes a slow and progressive DNA virus infection associated with epithelial remodeling and barrier disruption, whereas RVs induce rapid and transient RNA virus replication with limited tissue damage. Consistent with their respective viral life cycle, AdV accumulated progressively in our model over several days and was associated with basolateral viral leakage showing barrier disintegration, while both RV-A and RV-C peaked within 24 h and rapidly declined without detectable loss of epithelial integrity.

Unexpectedly, however, none of the three viruses displayed enhanced replication in pediatric BE compared to adult BE tested on multiple donors. This finding contrasts with clinical and epidemiological observations but is consistent with the fact that severe pediatric RV disease is frequently associated with underlying airway pathology, particularly asthma ^5,6^. Importantly, our cohort specifically excluded donors with severe chronic respiratory or immune disorders, suggesting that BE derived from healthy pediatric donors are not intrinsically more permissive to AdV or RV infection compared to adult BE. The absence of age-related differences in virus susceptibility coincided with morphological and physiological similarities. Adult and pediatric BE showed comparable epithelial organization, differentiation status, and barrier properties, suggesting that healthy BEs from both age groups largely share their intrinsic characteristics. Single-cell transcriptomic analysis further supported this conclusion, revealing highly comparable cellular compositions and transcriptional profiles, with no major differentially regulated pathways detected between both groups.

In our previous study using the same ALI-derived BE model during SARS-CoV-2 infection, we already observed that pediatric and adult epithelia displayed comparable epithelial organization and baseline permissiveness, while differences emerged primarily at the level of infection-induced antiviral responses, notably through enhanced type III interferon-associated restriction in a subset of pediatric donors ^1^. Together, these findings argue against the existence of major constitutive age-dependent epithelial determinants in healthy individuals. Instead, they suggest that age-related clinical susceptibility likely results from dynamic antiviral and inflammatory responses, pre-existing airway conditions, immune cell interactions, or infection history rather than from fundamentally distinct epithelial architectures or baseline permissiveness.

This conclusion was further supported by scRNA-seq analysis of infected epithelia. Indeed, no major age-dependent differences were observed between adult and pediatric BE following infection. In both age populations, AdV infected a broad range of cell types, including basal, suprabasal, and multiciliated cells, and induced the emergence of a distinct population of highly infected “AdV-specific” cells (>90% infected cells based on viral transcripts) characterized by a distinct expression signature. Interestingly, this population was uniquely associated with AdV infection and was not observed following RVA or RVC infection. In contrast, both RV species displayed a much more restricted tropism, primarily targeting multiciliated cells with only rare infection of goblet or basal cells. The conservation of viral tropism and infection efficiency across age groups further supports the idea that the clinical differences observed *in viv*o are unlikely caused by major intrinsic epithelial differences alone. Rather, as suggested by our previous SARS-CoV-2 study and others, age-dependent disease outcomes may depend more strongly on the strength of epithelial antiviral signaling, cytokine production, and interactions with surrounding immune and stromal cells within the airway environment ^18–21^ .

Using scRNA-seq, we were able to demonstrate that BE infection with AdV, RVA or RVC induces a shared epithelium-wide transcriptional response dominated by interferon-stimulated genes (ISGs). Notably, ISG upregulation was observed to a similar extend in both infected and bystander cells, while interferon gene expression (IFN-λ and IFN-β) was restricted to infected cells. This observed pattern supports a model in which infected cells produce interferons, which subsequently induce ISG expression in neighboring bystander cells, establishing a broad antiviral state across the BE. The absence of significant differences in ISG induction between adult and pediatric BE further reinforces the idea that central epithelial antiviral responses are conserved across age groups, at least in this in vitro model.

One of the key findings from our scRNA-seq analysis is that the BE mounted a relatively focused ISG response, confined to a highly conserved ISG signature across all epithelial cell types, despite the marked biological differences between the input viruses. Moreover, we show that the response was sufficient to induce a cross-protection against secondary viral infections, suggesting that the interferon-driven epithelial response establishes a prolonged protective state within the BE. Such cross-protection mechanisms have previously been proposed based on epidemiological observations. Indeed, Gopal et al, compared the prevalence of sequential viral infections in patients with previously documented positive versus negative swabs across multiple studies ^11^. Their meta-analysis revealed a reduced prevalence of secondary respiratory viral infections following an initial positive viral detection, leading the authors to propose the existence of a transient cross-protective state of the respiratory tract mediated by broad, non-specific innate immune mechanisms rather than virus-specific adaptive immunity. Our findings provide experimental support for this model and suggest that the BE itself may actively contribute to this transient antiviral protection through the establishment of an interferon-mediated tissue-wide antiviral state.

*In vivo*, one of the main mechanisms proposed for such a cross protection involves immune cells such as memory T-cells ^22–25^. However, increasing evidence also points toward an active role of the airway epithelium itself in establishing a transient antiviral state. Moore et al, showed that rhinovirus infection triggers expression of antiviral airway genes that lower the risk to subsequent SARS-CoV-2 infection ^26^. Likewise, repeated nasal sprays of interferon *vs.* placebo reduced the incidence and severity of colds in volunteers challenged with human rhinovirus^27^. Together, these studies support the notion that interferon-driven antiviral programs within the respiratory tract can transiently protect against secondary viral infections, independently of virus-specific adaptive immunity.

Here we demonstrate, using our physiological BE model, that up regulation of a defined subset of ISGs is sufficient to confer BE protection against a subsequent viral infection. This finding is particularly noteworthy, as it suggests that not all ISGs contribute equally to antiviral resistance; instead, a subset of ISGs, induced by different viruses, may be the prime drivers of the cross-protective state. The temporal discordance between interferon production (which peaked and declined rapidly) and cross-protection (which persisted longer) suggests that sustained ISG expression, rather than transient interferon bursts, may be critical for maintaining the refractory state. The conservation of this signature across viruses with distinct replication strategies and kinetic such as AdV and RVs and across age groups underscores its fundamental role in epithelial antiviral defense.

Our findings provide experimental support for this model. Furthermore, using viruses with very different replication strategies and kinetics, including the slowly replicating DNA virus AdV and the rapidly replicating RNA viruses RVA and RVC, we observed for both virus groups a relatively restricted and conserved ISG signature across epithelial cell types. In addition, the distinct infection dynamics uncovered an important temporal aspect of this response. While interferon production peaked rapidly and declined early after infection following individual virus replication kinetics, the elicited antiviral protection persisted substantially longer, suggesting that sustained ISG expression following transient interferon production is key to maintaining cross-protection. This temporal dissociation between virus induced interferon signaling and prolonged antiviral resistance further supports a model in which interferons primarily act as initiating signals that trigger a more durable ISG-mediated antiviral program within the epithelium. However, simply replacing the primary viral infection with interferon application shortened the cross-protection suggesting that continuous viral replication may be required. The conservation of this response across distinct viral families and across age groups nevertheless underscores its likely fundamental role in epithelial antiviral defense.

In order to further investigate the role of this subset of ISGs in cross-protection we performed a RNA-seq analysis on cross protected BE at different time points and identified genes significantly correlated with cross-protection. Surprisingly very few ISGs were highly correlated with the level of BE cross protection. Among the identified ISGs, IFI44L emerged as a key mediator, showing the strongest correlation between its induced expression and the observed level of cross-protection.

IFI44L is a well-described interferon-stimulated gene induced during viral infection ^28^ and has been reported to be upregulated following infection by several respiratory viruses, including SARS-CoV-2, influenza virus, and RSV ^29,30,31^. Although the precise underlying antiviral mechanism remains incompletely understood, previous studies suggest that IFI44L may participate in innate immune signaling through interactions with the JAK/STAT1 pathway ^32,33^.

Here, we demonstrate that IFI44L expression is robustly induced following infection with RVA, RVC, and AdV, despite major biological differences between these viruses. Importantly, IFI44L expression levels strongly correlated with the magnitude of epithelial cross-protection. Functional validation confirmed this association, demonstrating that heterologous IFI44L overexpression in BE significantly reduced RVA infection. Together, these findings identify IFI44L as a potential key effector of the epithelial antiviral state and suggest that specific ISGs, rather than the global interferon response itself, may represent critical determinants of broad antiviral protection in the airway epithelium.

Few trade-offs of this study should be acknowledged. First, our BE model does not incorporate immune cells and other components of the respiratory microenvironment that could contribute to antiviral defense and disease outcome *in vivo*. However, this reduced complexity allows a specific analysis of BE-intrinsic antiviral responses without confounding influences from other cellular compartments. This allowed us to uncover epithelial determinants of cross-protection, which could have been obscured in the more intricate context of in vivo studies In contrast, this also may partly explain the absence of major age-dependent differences in our model despite the well-documented clinical disparities in respiratory virus susceptibility and severity between children and adults. It is likely that other factors beyond the epithelium itself, including systemic immune responses, immune maturation, microbiome composition, or environmental exposures, may help shaping age-specific outcomes. In addition, our study specifically used “healthy” donors without severe chronic respiratory disease. It is therefore possible that age-dependent epithelial differences become more pronounced in pathological contexts such as asthma, where impaired interferon responses have previously been reported. Such a pivotal role of interferon in BE response to viral infection may partially account for the physiopathology of asthma in which IFN-λ production has been shown to be deficient during asthma exacerbation ^34,35^. Further studies using asthmatic or otherwise susceptible patient cohorts will be required to address this question. Second, our analysis focused on a side-by side comparison of three biologically and clinically distinct viruses including rapidly replicating RNA rhinoviruses and the slower replicating DNA adenovirus. All three are major respiratory pathogens especially in the pediatric setting. The comparative approach is a major strength as, to the best of our knowledge, it has never been conducted at this scale. This allowed us to identify conserved antiviral epithelial responses across markedly different viral lifecycles and infection kinetics. Expanding this approach in the future to additional respiratory pathogens, including RSV, influenza virus, or SARS-CoV-2, may help determine how broadly such interferon-mediated cross-protective states operate within the airway epithelium.

Taken together our results have important implications for understanding the interplay between respiratory viruses and the host epithelium. The demonstration of transient, cross-viral protection in BE suggests that primary infections may shape the susceptibility of the airway to subsequent pathogens, potentially explaining the observed patterns of viral co-infections in clinical settings. The identification of a conserved ISG signature associated with protection opens new avenues for therapeutic intervention and understanding disease severity. Targeting specific ISGs, or the pathways that regulate their expression, could offer a strategy to induce a broad, virus-agnostic antiviral state in the airway epithelium, potentially mitigating the severity of respiratory infections.

## Materials and Methods

### Study population

A total of 43 adults were prospectively recruited at the University Hospital of Bordeaux, France, after surgical resection from the TUBE Cohort. A total of 30 children were prospectively recruited at the Children University Hospital of Bordeaux, France, after fiberoptic bronchoscopy from the BIRDIES or VIRCHILLD Cohort. All patients, adults and pediatric were non-asthmatics and with a normal to sub-normal lung function. All subjects gave their written informed consent to participate in the study after the nature of the procedure had been fully explained. All samples were deidentified prior to their use in this study.

### Cell culture

The bronchial epithelial cell culture was established from bronchial brushings or lung resection performed between the third and fifth bronchial generation from patients undergoing either elective surgery or fiberoptic bronchoscopy. Bronchial brushings or dissected epithelial explants were cultured using PneumaCult-Ex® expansion medium (StemCell Technologies) at 37 °C in 5% CO2 in 6 well plate. Basal cells were then transferred to a 75 cm² culture flask and cultured in PneumaCult-Ex Plus® medium in order to obtain sufficient numbers of basal cells. Then, 10^5^ basal cells were transfer on Transwell® polyester membranes (0.4 µm pore size) in PneumaCult-Ex Plus® medium. The following day, medium was removed, and inserts were placed in air–liquid interface using PneumaCult-ALI® medium (Stemcell) at basal level and allowed to differentiate for a minimum of 21 days at 37 °C with 5% CO2 to obtain a reconstituted and fully differentiated bronchial epithelium. Such a complete differentiation was assessed under a light microscope to assess ciliary beating and mucus production.

### Virus production

#### Production of RV-A-GFP and RV-C

The cDNA pA16-GFP plasmid encoding RVA-GFP, which contains the rhinovirus and GFP sequence and downstream a T7 promoter, and the cDNA pC15 encoding RVC, is a kind gift from Yury A Bochkov James Gern ^36^. The reverse genetics technique developed by Bochkov et al ^36^ was used to produce functional RV16-GFP and RVC virions. Briefly, for the initial production, the pA16-GFP or pC15 plasmid were linearized by incubation with respectively the restriction enzyme Eco53kI (Eco53kI, Thermo Scientific) at 10 U/μL or BstBI (BstBI, Thermo Scientific) at 10 U/μL. After purification the linearized DNA was transcribed in vitro into RNA by T7 polymerase kit (Promega) and then transfected into HeLa cells using lipofectamine (Invitrogen). 24 hours after transfection, the produced virus was purified, aliquoted and stored at -80°C. Titration of the virus was performed by digital PCR after extraction and reverse transcription (qPCR Transcriptomic platform, University of Bordeaux). Digital PCRs were prepared with the required reagent, QX200 ddPCR EvaGreen Supermix (Bio-Rad).

#### Production of RV-A

The human rhinovirus RV-A was purchased from ATCC, aliquoted and stored at -80°C.

#### Production and purification of AdV

Adenovirus 5 (HAdV-C5) was obtained from ATCC and first amplified in HEK293-αvβ5 cells. Virus was extracted from infected cells by repeated freeze-thaw cycles and purified via double CsCL gradient banding as previously described ^37^. Concentrated virus was then dialyzed against PBS/10% glycerol in Slide-A-Lyser Pierce dialysis cassette, aliquoted and stored at -80℃. Virus concentration was evaluated by measuring OD (optical density) at a wavelength 260 nm.

### Virus infection (RVA, RVC, AdV and RVA-GFP)

Apical sides of the BE cells were infected with 100µl of DMEM medium with either AdV, RV-A, RV-A GFP and RV-C during 1 hour then medium was completely removed. Optimal infective doses were determined empirically. AdV was applied at high MOI (2.3×10^9^ physical particles per epithelium) to achieve robust and reproducible infection. Both RVs were applied at 0.1 MOI ^38^.

### Immunostaining

BE inserts were fixed in 4% PFA for 30 minutes (500 µl applied to the basolateral side, 200 µl applied to apical side). Fixed BE inserts were permeabilized with 0.5 % Triton X100 (in 1 X PBS) for 10 min at room temperature, washed and blocked in IF buffer containing PBS with 10% fetal bovine serum and 0,05% of saponin for 1h. Primary antibodies were diluted in IF buffer and applied to the BE overnight at 4℃. Immunostaining was performed using antibodies against alpha-acetylated tubulin to label multiciliated cells ([EPR16772]- Abcam), anti-MUC5AC to label goblet cells ([EPR16904], Abcam) and anti-cytokeratin 5 to label basal cells ([EP1601Y], Abcam). Adenovirus infected cells were detected using antibodies against DBP labeling replication center (kindly provided by R. Iggo, Bordeaux, France). Rhinovirus A and C infected cells were detected with antibodies against capsid protein VP3 (G47A, Invitrogen). Secondary antibodies were diluted in IF buffer and incubated with the BE for 2h at room temperature. Nuclei or actin cytoskeleton was stained with DAPI or Phalloidin (Invitrogen, A30104) at 1/400 dilution of stock concentration (*i.e* 3mM and 66µM respectively).

### Imaging and live-imaging

#### Image Acquisition

Confocal imaging of stained BE inserts was performed at the Bordeaux Imaging Center (BIC) using a Leica DMI8 inverted microscope (Leica Microsystems, Wetzlar, Germany) equipped with a Photometrics Prime 95B sCMOS camera (Photometrics, Tucson, AZ, USA). The microscope was fitted with a 100X oil-immersion objective (HC PL APO 100×/1.40 NA, Leica Microsystems). Fluorescence excitation was provided by a multi-line diode laser system (405 nm, 491 nm, 561 nm, and 642 nm). Immersion oil was applied to ensure optimal resolution and signal-to-noise ratio. To monitor RVA GFP infection in living BE, whole BE inserts were mounted on a motorized stage using a Leica DMI6000 inverted microscope (Leica Microsystems, Wetzlar, Germany) equipped with environmental chamber equilibrated at 37°C and a Hamamatsu Orca Flash4.0 sCMOS camera (Hamamatsu Photonics, Hamamatsu, Japan). The microscopes and cameras were controlled via MetaMorph software (version 7.10.6, Molecular Devices, San Jose, CA, USA). BEs were automatically imaged using a 20x objective using the ScanSlide plugin (Molecular Devices) to automate the acquisition of tiled images across the entire sample. Tiling parameters, including overlap percentage (10%), focus settings, and exposure time, were optimized to ensure seamless coverage of the epithelial layer.

#### Image Reconstruction and Analysis

Tiled images were stitched into a single high-resolution mosaic using a custom ImageJ macro. The stitching process aligned individual tiles based on overlapping regions, correcting for minor shifts in the X, Y, and Z axes. The resulting stitched images were then analyzed for quantitative measurements, such as cell density, morphology, or fluorescence intensity, using ImageJ’s built-in tools or additional plugins as required (The ImageJ macro used for stitching and analysis is available upon request).

### Transepithelial resistance (TEER)

BE permeability was obtained by measuring the transepithelial resistance using EVOM 3 device with STX4 electrode (EVOM, World precision instruments) following the manufacturer’s recommendations.

### Ciliary beating frequency

Ciliary beating frequency of BEs were measured using videomicroscopy Leica DMi8 (Leica Microsystems) coupled to a high-speed camera sCMOS Flash 4.0 camera (Hamamatsu), available at The Bordeaux Imaging Center. The illumination system used was Cool LED PE-4000 (CoolLED) associated to a HCX PL Fluotar L 40X dry 0.6 NA PH2 objective. BEs were placed in a 37°C chamber using an incubator box equipped with a gas heating system (Pecon GmbH). The acquisitions were carried out during 2 seconds at a frequency of 1000 images/sec so a total of 2001 images, on a defined zone (100×100 pixels) using MetaMorph software (Molecular Devices). Using MATLAB program, the images were analyzed after application of the Fourier transform to determine the most represented beat frequency ^39^.

### RNA extraction and Reverse transcription

Total BE RNA was extracted from cells using a Qiagen RNeasy kit (Qiagen) by following the manufacturer’s recommendations. Reverse transcription was perform using iScript Reverse Transcription Supermix (BioRad) by following the manufacturer’s recommendations.

### TaqMan PCR (*RV-A* and *RV-C*)

After RNA extraction and Reverse transcription, RV quantification was performed using iTaq Universal Probes Supermix (BioRad) by following the manufacturer’s recommendations. (Human rhinovirus reverse primer 5’AAACACGGACACCCAAAGTAGT-3’, Human rhinovirus forward primer 5’-AGCCTGCGTGGCKGCC-3’, Human rhinovirus probe 5’FAM-CTCCGGCCCCTGAAGGCTAA-3’.

### Real-time qPCR (*AdV*)

SybrGreen staining and the Adenovirus 5 hexon primers (AQ1 and AQ2) were used for qPCR reaction. The sequence of primers was obtained from [*Heim et.al, J Med Virol., 2003*] to amplify *hexon* gene conserved among human Adenovirus species A-F.

The primer sequences are as follows: *5’ Hexon AQ2* - 5’-GCC-CCA-GTG-GTC-TTA-CAT-GCA-CAT-C-3’, *3’ Hexon AQ1*- 5’-GCC-ACG-GTG-GGG-TTT-CTA-AAC-TT-3’.

### RT-PCR ISG

RT-qPCR for relative ISG expression was performed using Perfecta Sybergreen (Quanta bio) following manufacturer protocol. The primers used for this RT-qPCR are listed in the table below:

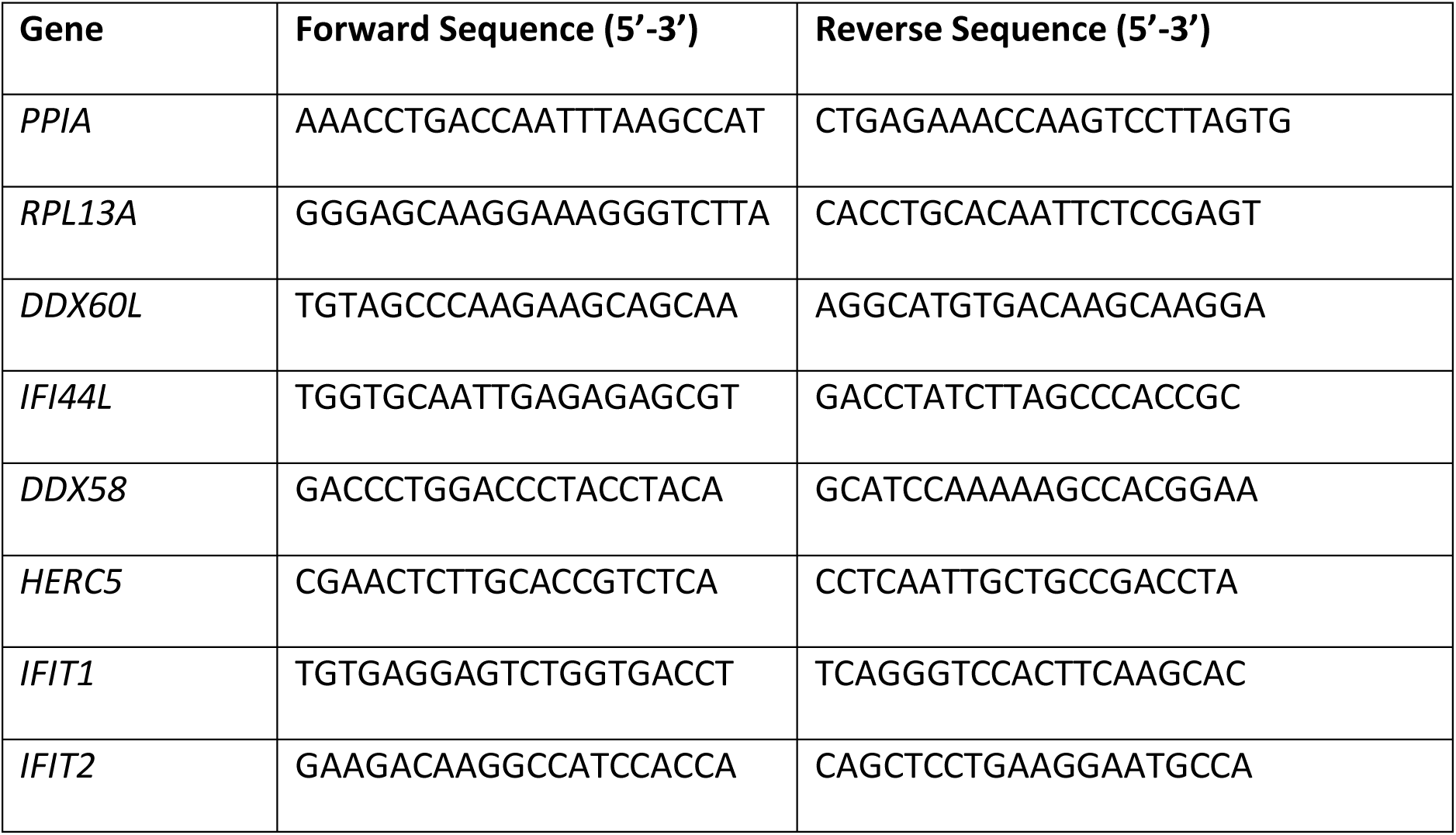
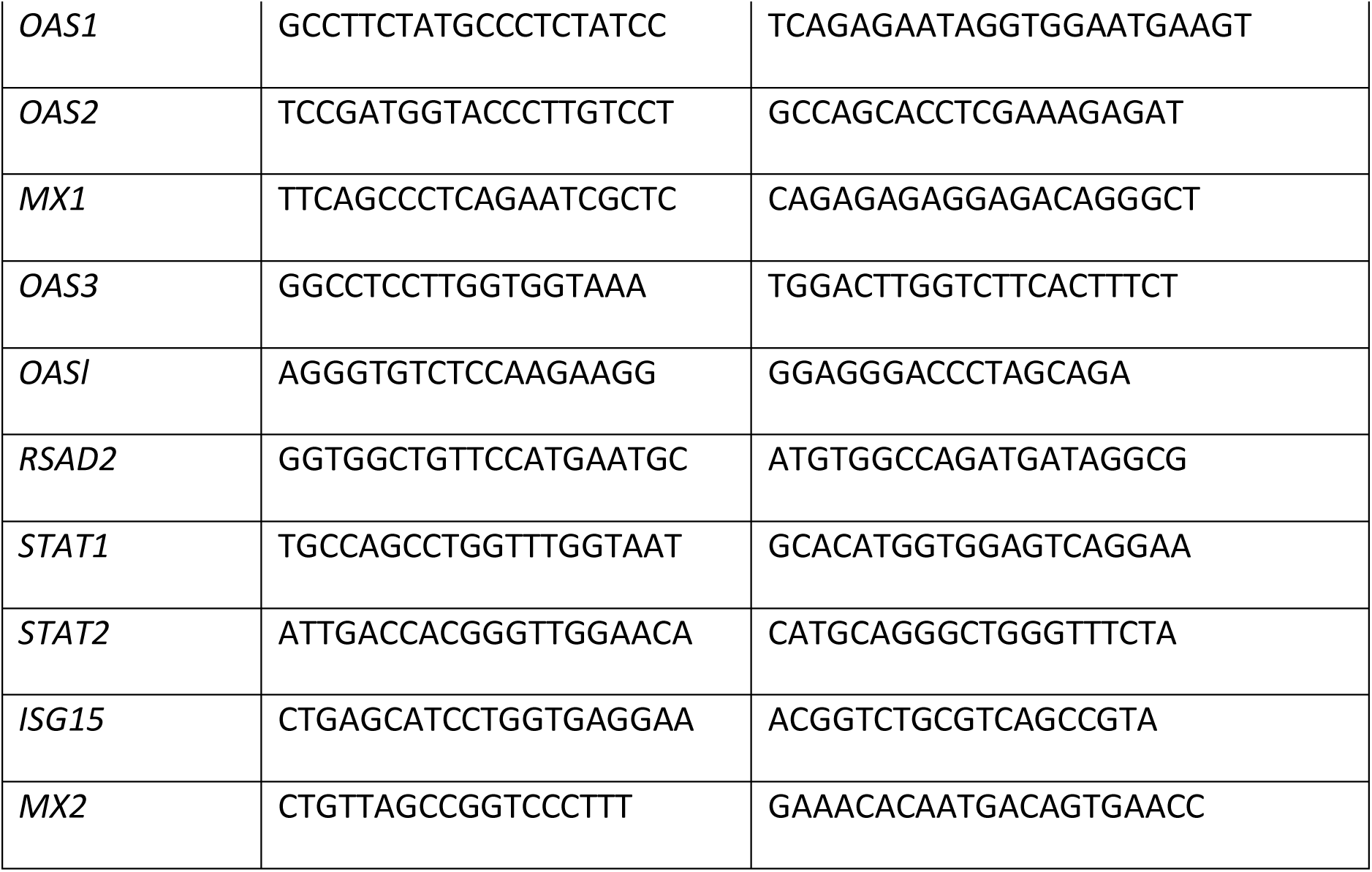

### ELISA

Interferon response (IFN β and IFN λ1-3) were appreciated using ELISA (Enzyme-Linked Immuno-Sorbent Assay). BE cell culture media were immediately stored and frozen at -80°C for optimal preservation. ELISA for interferons (Bio-Techne) were performed according to the manufacturer’s recommendation and measured at 450 nm using the SPECTROstar device and the associated SPECTROstar Nano MARS software.

### Lentivirus production

Lentiviral vector production was done by the service platform for lentiviral vector production “Vect’UB” of the TMBCore UAR3427/US05, Bordeaux University. Lentiviral vector was produced by transient transfection of 293T cells according to standard protocols. In brief, subconfluent 293T cells were co-transfected with lentiviral genome (psPAX2) ^40^ with an envelope coding plasmid (pMD2G-VSVG) and vector constructs expressing IFI44L mRNA and mCherry under MNDU3 promoter (vector builder) by calcium phosphate precipitation. LVs were harvested 48 hours post-transfection and concentrated by ultrafiltration columns. Viral titers of pLV lentivectors were determined by transducing 293T cells with serial dilutions of viral supernatant and integrated provirus copy number was quantified 5 days later by qPCR method.

### Western blotting

Total BE lysis was performed using a RIPA lysis buffer (Sigma-Aldrich). Total cellular extracts were loaded onto a 4-20 % SDS-PAGE gel (Bio-Rad), transferred onto nitrocellulose membrane. Antibody directed against IFI44L (Abcam) was used. HRP-coupled secondary antibodies were used for revelation using ChemiDoc imaging instrument (Bio-Rad). Protein expressions were normalized using total loading protein intensity (Stain-free system Bio-Rad).

### Single-cell RNA-seq

#### Cell dissociation

To perform single-cell analysis, cells differentiated on Transwells® were harvested by incubating with 0.1% protease type XIV Streptomyces griseus in HBSS for 4-6 hours at 4°C, until dissociation was clearly visible. Then, cells were gently detached from the Transwell® by pipetting and transferred into a microtube. An excess of HBSS containing 2% BSA was added. Cells were centrifuged at 150 g for 5 min and resuspended in 500 µL of supplemented HBSS containing 2% BSA, centrifuged again at 150 g for 5 min and resuspended in 500 µL HBSS. Finally, cell suspensions were filtered through a 40 µm porosity Flowmi^TM^ Cell Strainer (Bel-Art), centrifuged at 150 for 5 min and resuspended in 150 µL of HBSS. Cell concentration and viability were measured with a Luna automated cell counter (Logos Biosystems), after incubation with NucGreen to detect dead cells (Thermo Fisher scientific). All steps were performed on ice. Cells were fixed and then barcoded according to the manufacturer’s protocol (Parse Biosciences Evercode^TM^ WT v2) to obtain single cell 3’ libraries for Illumina sequencing. The libraries were sequenced with a NextSeq 2000 P3 Reagents (100 cycles) in paired-end mode at length of 28 bases for R1 and 90 bases for R2 with 2 index reads of 10 bases. Raw sequencing data were processed using the split-pipe pipeline (version 1.0.3p) from Parse Biosciences-Pipeline, with default parameters, with sample list barcoding provided in Supplementary Table S2A and S2B and aligned to the GRCh38 human reference genome (Ensembl release 108). Viral genomes, including Adenovirus (ADV), Rhinovirus A (RVA), and Rhinovirus C (RVA), were incorporated into the reference in order to quantify reads mapping to viral sequence.

#### Single-cell data analysis

##### Preprocessing, integration, normalization and clustering

Individual dataset analysis was performed using Seurat standard analysis pipeline v4.3.0 ^41^. For each sample, cells were filtered out based on number of expressed features, dropout percentage, library size and mitochondrial gene percentage. Thresholds were selected by visually inspecting violin plots in order to remove the most extreme outliers (for samples details and filtering see Supplementary Table 3). Then, we split the data into 2 datasets, with the control samples in a first dataset in order to compare adult versus pediatric samples and a second dataset with control, ADV, RVA and RVC samples to compare virus effected regardless the age. We performed integration with Seurat integration and we constructed a k-nearest neighbour graph with default parameters. The Uniform Manifold Approximation and Projection (UMAP) representation was used for visualization of integrations into two dimensions. Cell-level normalization was performed using NormalizeData function. Highly variable genes were selected for following analysis based on their expression level and variance. Clustering was performed with a resolution parameter of 0.5 and using Seurat function. Differential analysis was again performed using Seurat FindAllMarkers and FindMarkers functions based on non-parametric Wilcoxon rank sum test.

##### Cell type annotation

We used FindAllMarkers function to identify gene markers for each cluster found previously. We then annotated on the basis of these gene markers and by visualizing the expression of our gene markers.

##### Infected versus bystander cell classification

To identify the cellular compartments that were most frequently targeted by each virus and to assess their impact across cell types, we classified cells into infected and bystander populations for each viral condition (ADV, RVA, and RVC) at the single-sample level based on viral transcript abundance. This classification was performed independently for each virus, enabling downstream differential expression analyses between infected and bystander cells to identify host genes associated with viral infection status and to determine which cell types are more strongly affected by each virus. Cells were classified as “infected” if the number of normalized viral reads exceeded a virus-specific threshold. These thresholds were defined empirically by evaluating a range of thresholds from 0 to 5 with a step size of 0.1. The final threshold was selected to ensure that the false positive rate in control (uninfected) samples remained below 5%, in order to account for background signal and technical noise. The resulting cut-offs were 1.8 for ADV, 1 for RVA, and 0 for RVC (normalized counts).

Cells with normalized viral read counts below these thresholds were classified as “bystander” cells.

##### Analysis of cell type proportions

We used the propeller function (with asin transformation) from speckle package (v0.0.3) ^42^ to compare cell type proportion between age (adult vs. child) in non-infected primary BE. Supplementary Table S4 shows the detailed cell type proportions.

##### Differential Expression

We performed differential expression analysis by using the run_de function from Libra package (v1.0.0) (https://github.com/neurorestore/Libra) and using pseudobulk method with edgeR.

### Bulk RNA-sequencing

#### RNA-seq library preparation

RNA was extracted as detailed above. RNA sequencing was performed by Genewiz (Azenta Life Sciences). Libraries were prepared using a standard RNA-seq protocol with poly(A) selection and sequenced on an Illumina NovaSeq X Plus platform according to the manufacturer’s standard procedures.

#### RNA-seq Data Processing

Raw sequencing reads (FASTQ files) were aligned to the human reference genome (GRCh38, Ensembl release 108) using STAR (v2.7.10a) ^43^. Viral genomes, including Adenovirus, Rhinovirus A, and Rhinovirus C, were incorporated into the reference to enable quantification of reads mapping to viral sequences. Alignment was performed with the following parameters:

--readFilesCommand zcat --runThreadN 20 --outFilterType BySJout –outFilterMultimapNmax 20 --alignSJoverhangMin 8 --alignSJDBoverhangMin 1 --outFilterMismatchNmax 999 -- outFilterMismatchNoverLmax 0.04 --alignIntronMin 20 --alignIntronMa× 1000000 --alignMatesGapMa× 1000000 --outReadsUnmapped Fastx.

The resulting BAM files were sorted and indexed using SAMtools (v1.15.1) ^44^. Then, gene-level quantification was performed using featureCounts (v2.0.3) ^45^ with the following parameters: -g gene_id -t exon --primary -p --countReadPairs -B -C -s 0 -T 20

#### Differential Gene Expression

We performed differential gene expression paired analysis across time points (Day 1, Day 5, and Day 12) to compare the effect of repeated Rhinovirus infection on the various viruses initially present (Adenovirus, Rhinovirus A, Rhinovirus C) with uninfected controls. We used DESEq2 ^46^ paired comparison and genes were retained for analysis if they met the following thresholds: a minimum of 5 counts (nbCountsMin = 5) in at least 3 samples (nbSamplesMin = 3). Pairwise comparisons were conducted to identify genes differentially expressed between time points. To assess associations with the continuous variable “Cross_protection_GFP_Control”, each virus condition was compared independently to the control donor using limma-voom package ^17,47^. We transformed the “Cross_protection_GFP_Control” by normalizing it by its -log10 resulting to the “Cross_protection_neg_log10”.

#### Cell type proportions

Cell type proportions in the bulk RNA-seq samples were estimated using AutoGeneS in Python (v1.0.4), using the single-cell virus control dataset as a reference. ADV-specific cell populations, present in only one experimental condition, were excluded from the reference to avoid introducing bias into the marker gene selection. The top 4,000 highly variable genes were used as input features, and the genetic algorithm (ngen=500, offspring_size=100) was run to select an optimal set of 400 marker genes (mode=’fixed’), which were subsequently used to deconvolve bulk sample proportions via a Nu-Support Vector Regression (NuSVR) model. The resulting proportions were arcsine square-root transformed, following the propeller approach, and compared across viral conditions and time points using limma (v3.46.0), with duplicateCorrelation to account for repeated measures across shared donors. Specific contrasts (virus vs. control at each day, and between consecutive time points) were tested, with Benjamini-Hochberg correction for multiple testing.

Metadata associated with the RNA-seq dataset are provided in Supplementary Table S5.

## Data availability

Single-cell RNA-seq data is available under GEO and EGA platfoms where data can be browsed.

## AUTHORS CONTRIBUTIONS

TA performed the vast majority of the *in vitro* experiments and wrote the first draft of the manuscript GP, GS, BSR, FM, RB, RF and LN technically assisted with the *in vitro* experiments. BP and FB conducted patient recruitment. FM, GP and LEZ performed the Single cell and RNAseq experiments and analysis. BP, BP and EP helped to design the study and edited the manuscript. ZLE, HW and TT conceived the project, designed and supervised the study, analyzed the data, and edited the manuscript.

## ACKNOWLEDGEMENTS

We thank the study participants and the staff of the Thoracic Surgery, Pathology, Respiratory, Lung Function Testing departments from the University Hospital of Bordeaux (Bordeaux, France), All members from the adults and pediatric clinical investigation center of the University Hospital of Bordeaux (Bordeaux, France). Microscopy was performed at BIC, a service unit of the CNRS-INSERM and Bordeaux University, a member of the national BioImaging infrastructure of France supported by the French National Research Agency (ANR-10-INBS-04). Single cell sequencing was performed by the UCA GenomiX platform, with supports from the France 2030 program (Respirera : ANR-23-IAHU-0007, 4D-OMICs : ANR-21-ESRE-0052, 3IA : ANR-19-P3IA-0002; France Génomique: ANR-10-INBS-09-03) and by Conseil départemental 06.

## Supplemental Figure legends

**Figure S1.**
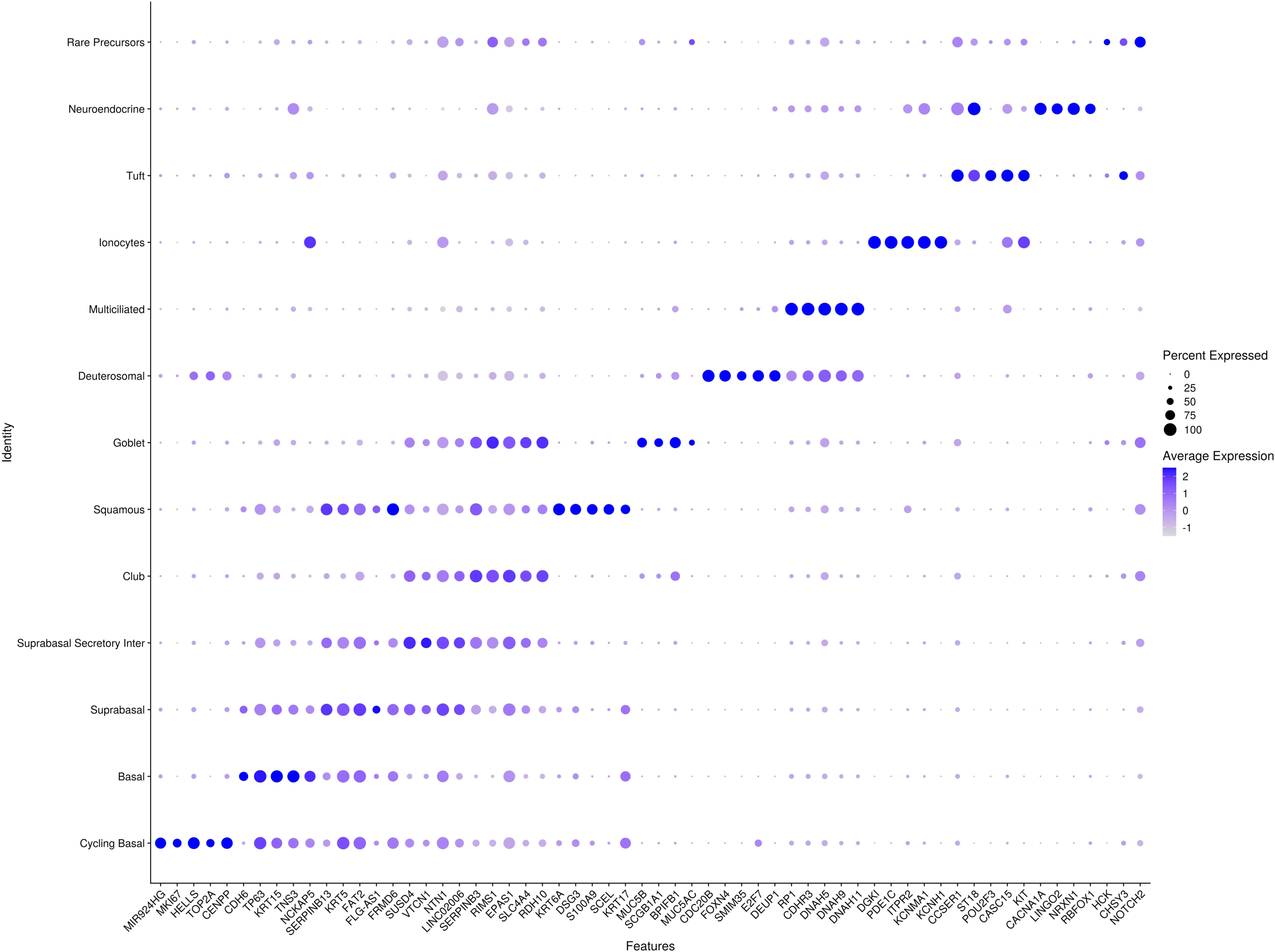
Gene expression signature of epithelial cell subsets in non-infected BE. All donors merged (n= 3 adults and 3 pediatric). The size of the dots corresponds to the percent expressed, and color indicate average expression.

**Figure S2.**
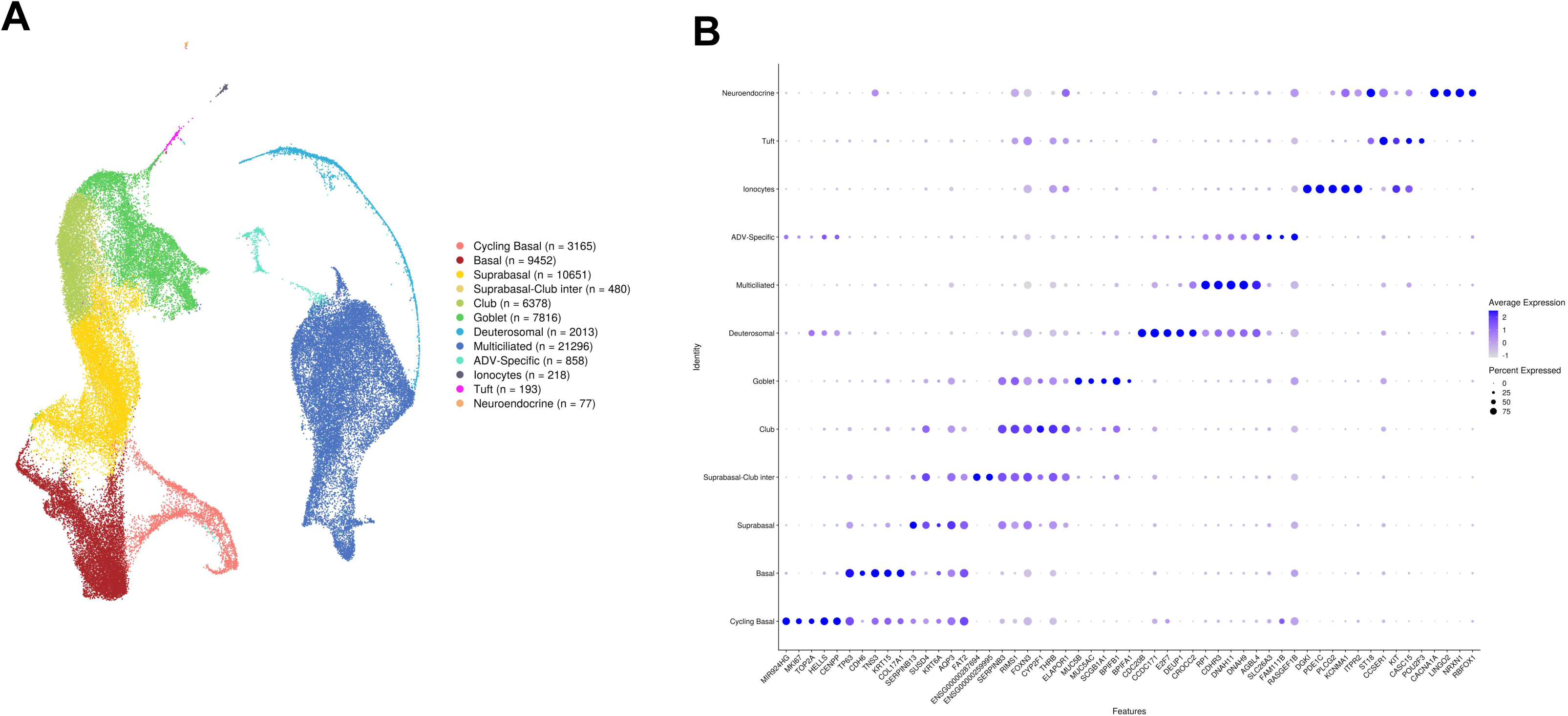
**A.** UMAP projection of the different cell types. The feature plots display the subset of airway epithelial cells obtained from infected adult and pediatric primary BE (n= 3 adults and 3 pediatric). **B**. Gene expression signature of epithelial cell subsets in infected BE. All donors merged (n= 3 adults and 3 pediatric). The size of the dots corresponds to the percent expressed, and color indicate average expression.

**Figure S3.**
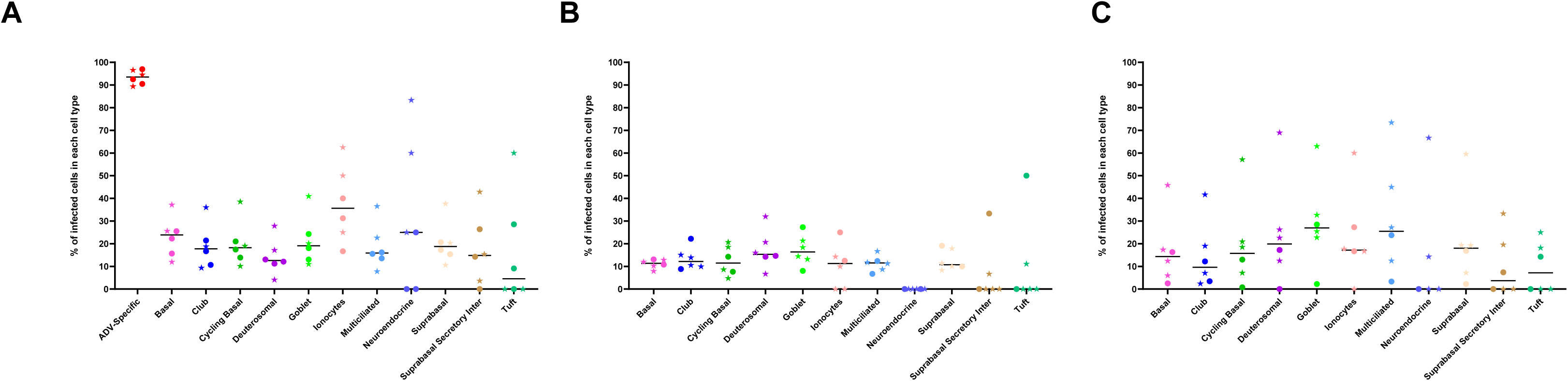
Percentage of AdV (**A**), RVA (**B**) and RVC (**C**) infected cells within each cell type in adult (circles) and pediatric (stars) primary BE.

**Figure S4.**
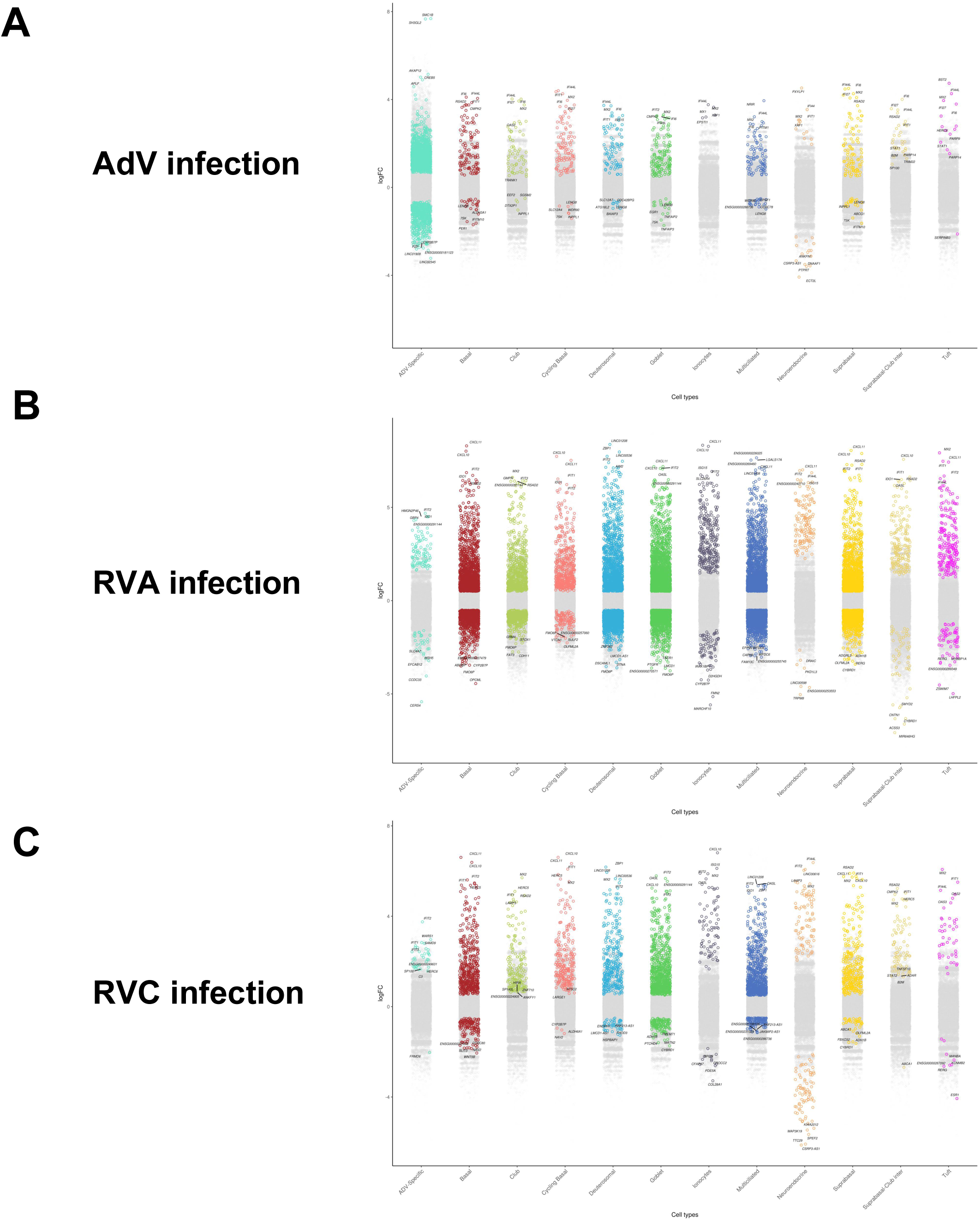
Strip plots of differential genes expression (log fold change) in each cell types following infection with either AdV **(A)**, RVA **(B)** or RVC **(C)** infection (n= 3 adults and 3 pediatric).

**Figure S5.**
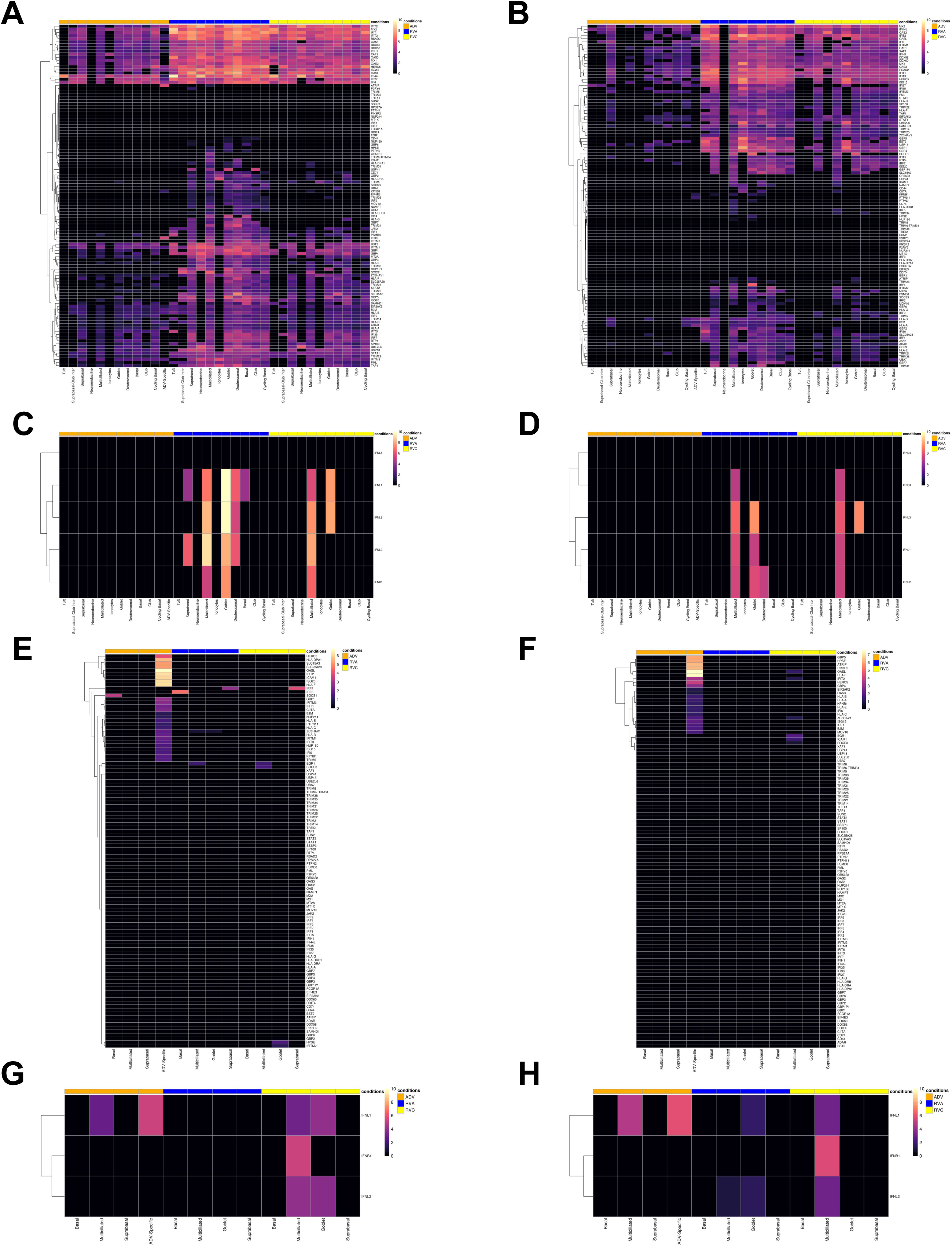
**A-B.** Heatmap of clustered Interferon induced genes upregulation for each cell type in adult only (n= 3) **(A)** or pediatric only (n= 3) **(B)** BE. ISGs activation was assessed between AdV, RVA or RVC infected primary BE compared to non-infected BE. **C-D** Heatmap of clustered Interferon genes upregulation for each cell type in adult only **(C)** or pediatric only **(D)** BE. Interferon genes activation was assessed between AdV, RVA or RVC infected primary BE compared to non-infected BE. **E-F.** Heatmap of clustered Interferon induced genes upregulation in cell type targeted by either AdV, RVA or RVC in adult only **(E)** or pediatric only **(F)** BE. ISGs activation was assessed between AdV, RVA or RVC infected cells and bystander non-infected cells from the same BE. **G-H.** Heatmap of clustered Interferon genes upregulation in cell type targeted by either AdV, RVA or RVC in adult only **(G)** or pediatric only **(H)** BE. Interferon genes activation was assessed between AdV, RVA or RVC infected cells and bystander non-infected cells from the same BE.

**Figure S6.**
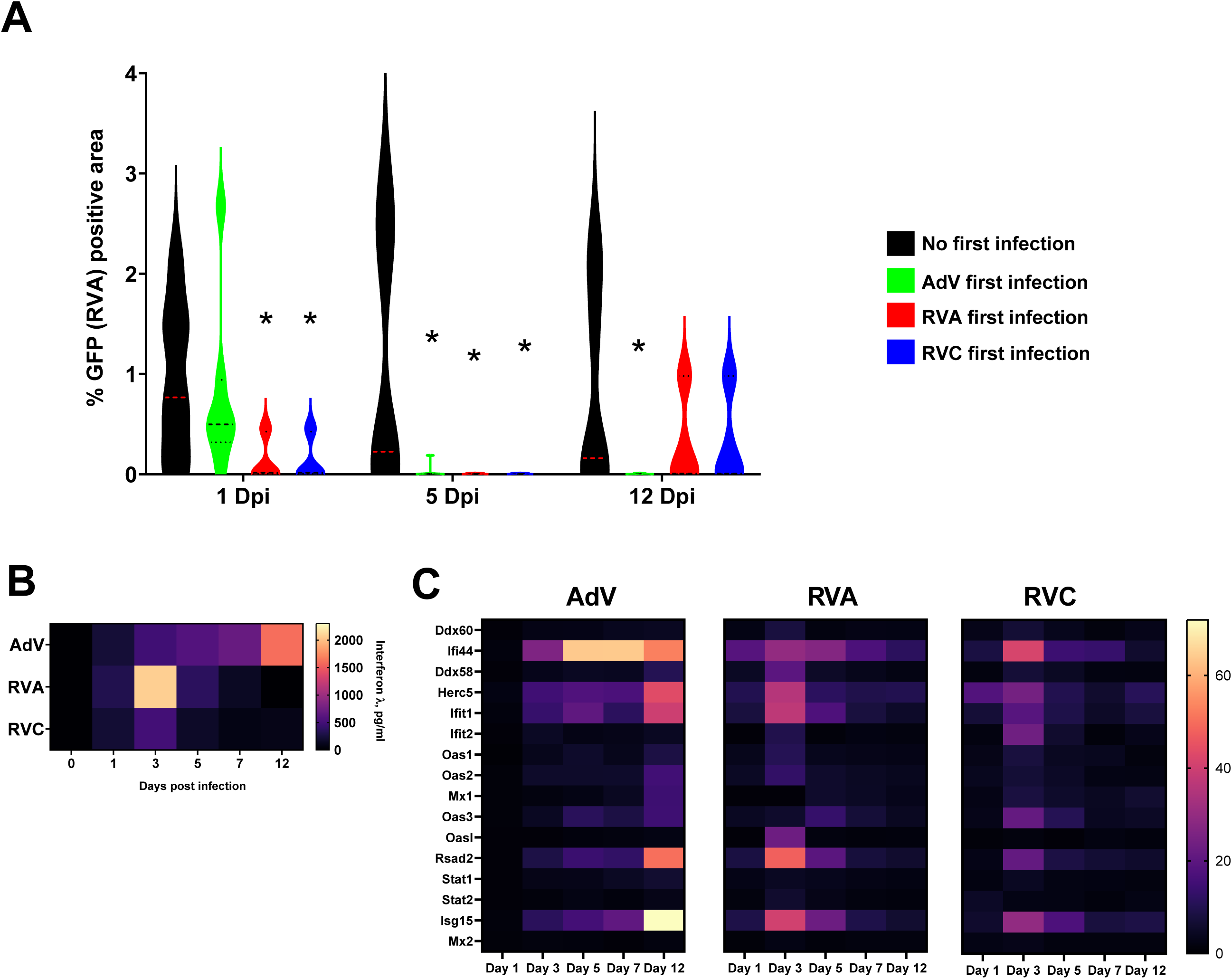
**A.** Percentage of GFP positive area induced by RVA-GFP infection in AdV (green), RVA (red), RVC (blue) and non-infected control (black) pre-infected pediatric primary BE after 1-, 5- and 12-day post infection (n=7). **B.** Heatmap of the kinetics of interferon λ release in AdV, RVA or RVC infected pediatric primary BE assessed by ELISA (n=9). **C.** Heatmap of the kinetics of ISGs fold increase mRNA expression in AdV, RVA and RVC infected pediatric primary BE assess by real time PCR (n=7).

**Figure S7.**
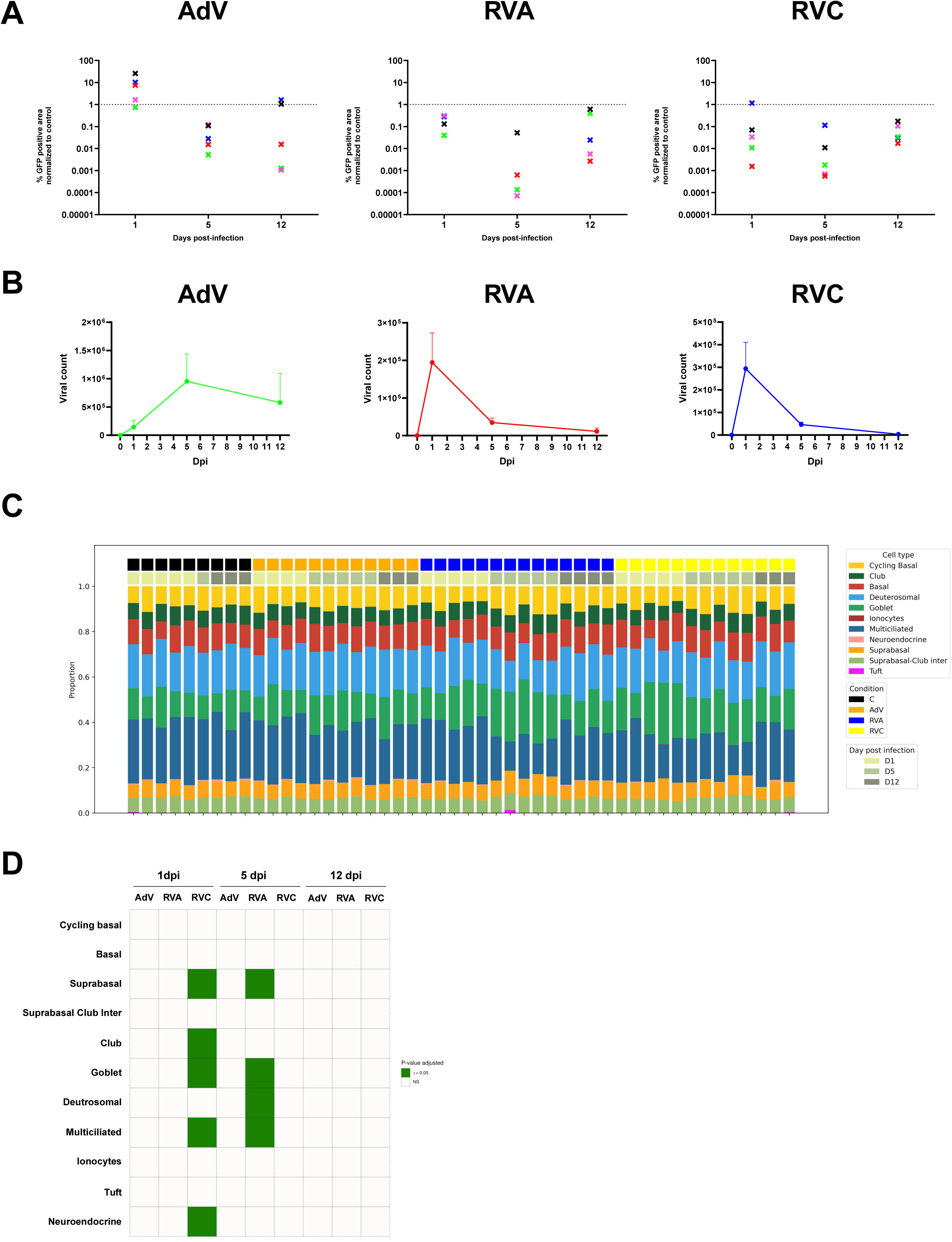
**A.** Percentage of GFP positive area (induced by RVA-GFP infection) in either AdV, RVA or RVC first infected BE after 1, 5 or 12 dpi. Data are presented as normalized to non-pre-infected control GFP positive area. Each value represents one sample and colors represent samples from the same patient at different time point. **B.** Viral counts in AdV, RVA and RVC infected adult primary BE after 1,5 and 12 dpi. **C.** Proportions of each cell type in all sample (AdV, RVA, RVC-infected and non-infected at 1, 5 and 12 dpi) obtain using cell deconvolution from RNA-seq data.

**Figure S8.**
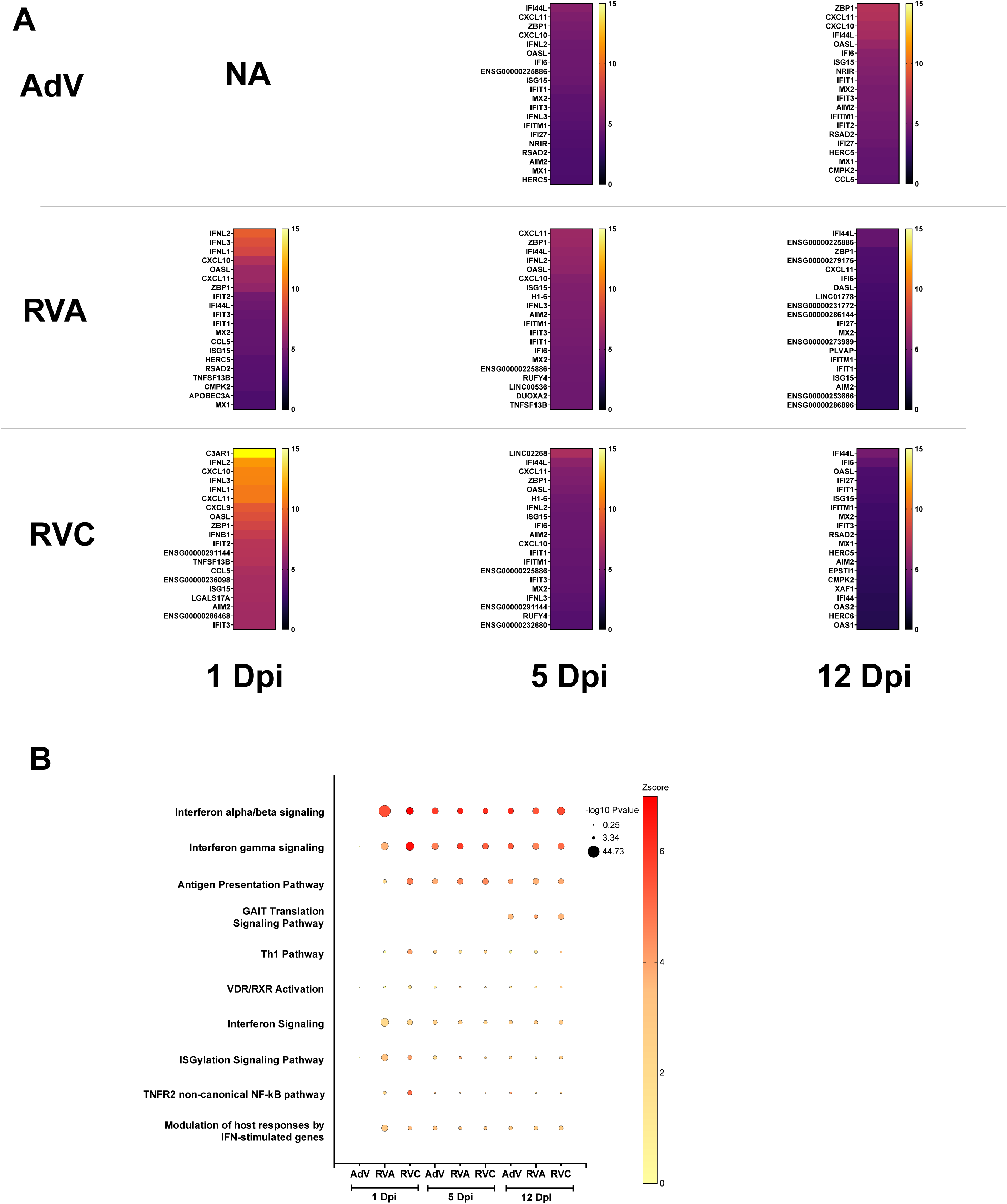
**A.** Heatmap of the top 20 significant overexpressed genes after AdV, RVA or RVC infection at 1, 5 or 12 dpi. **B.** Signaling pathways enrichment analyses were performed with human gene names using IPA platform. Comparison was performed between each BE infected condition (AdV, RVA and RVC) and non-infected BE at 1, 5 and 12 dpi. The size of the dots corresponds to the p-values, and color indicate the value of the z-score.

**Figure S9.**
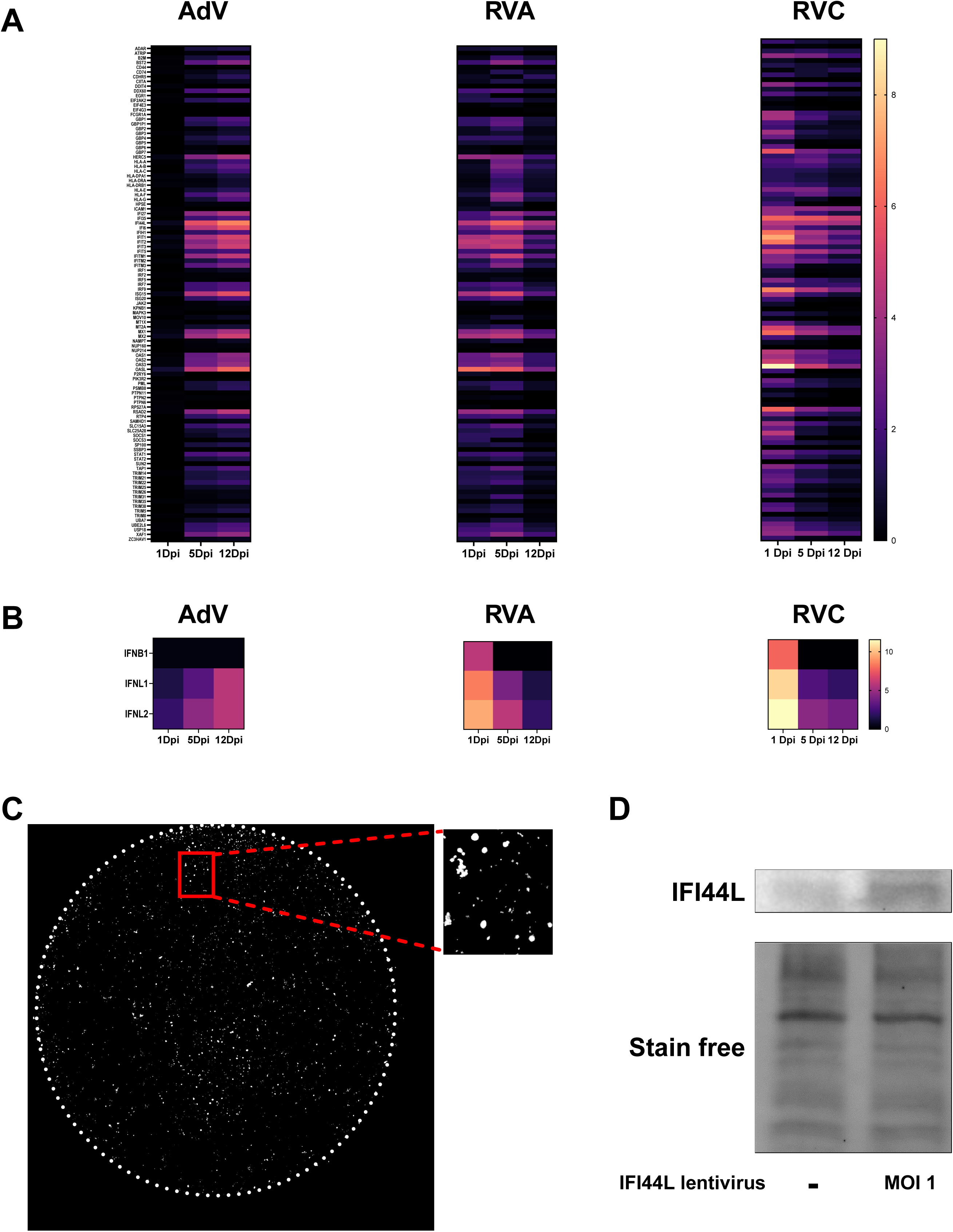
**A.** Heatmap of the kinetic of Interferon induced genes upregulation (assess by RNA-seq) in BE infected with either AdV, RVA or RVC after 1, 5 or 12-dpi. Interferon induced genes activation was assessed between AdV, RVA or RVC infected cells and corresponding non-infected cells from the same BE at the same time point. **B.** Heatmap of the kinetic of Interferon genes upregulation (assess by RNA-seq) in BE infected with either AdV, RVA or RVC after 1, 5 or 12-dpi. Interferon genes activation was assessed between AdV, RVA or RVC infected cells and corresponding non-infected cells from the same BE at the same time point. **C.** Representative images of mCherry signal induced by lentivirus transduction (magenta) in whole BE. **D.** Western blot analysis of IFI44L expression in BE transduced by IFI44L expressing lentivirus compare to non-transduced BE.

---

**Supplementary Table S1.**
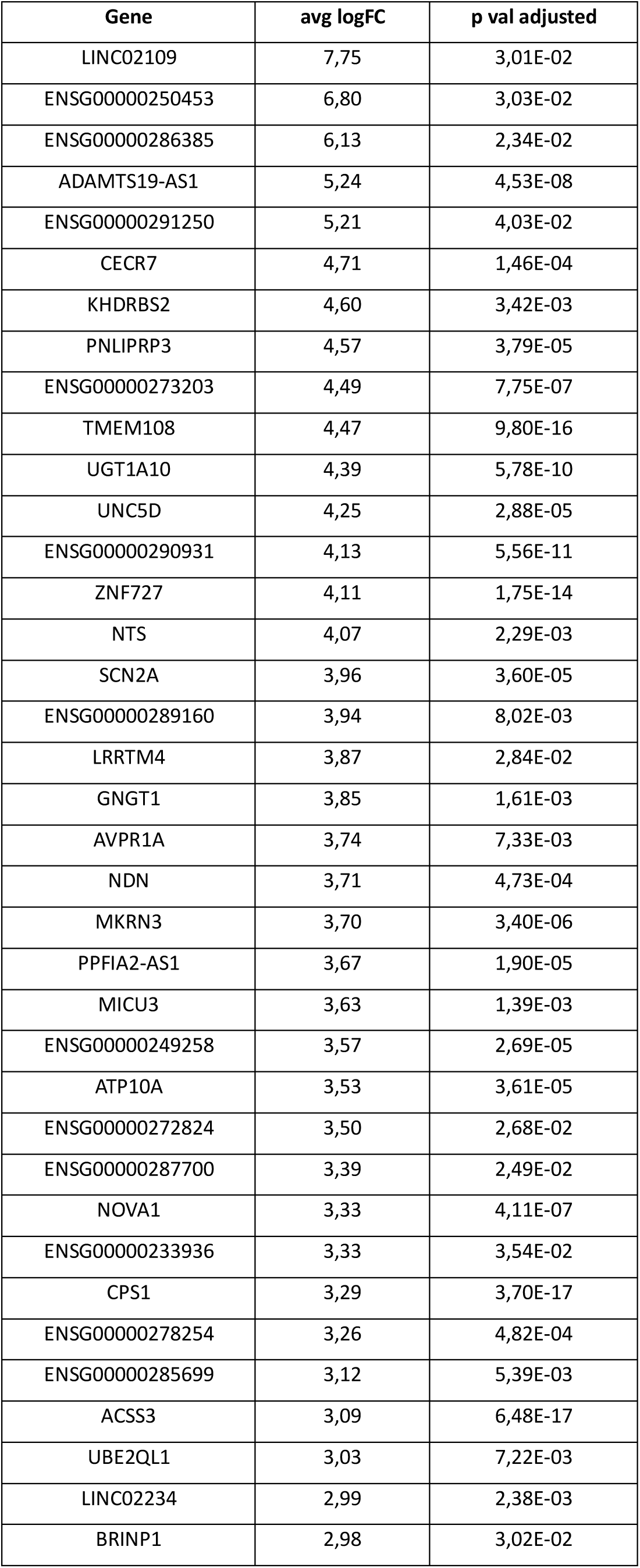
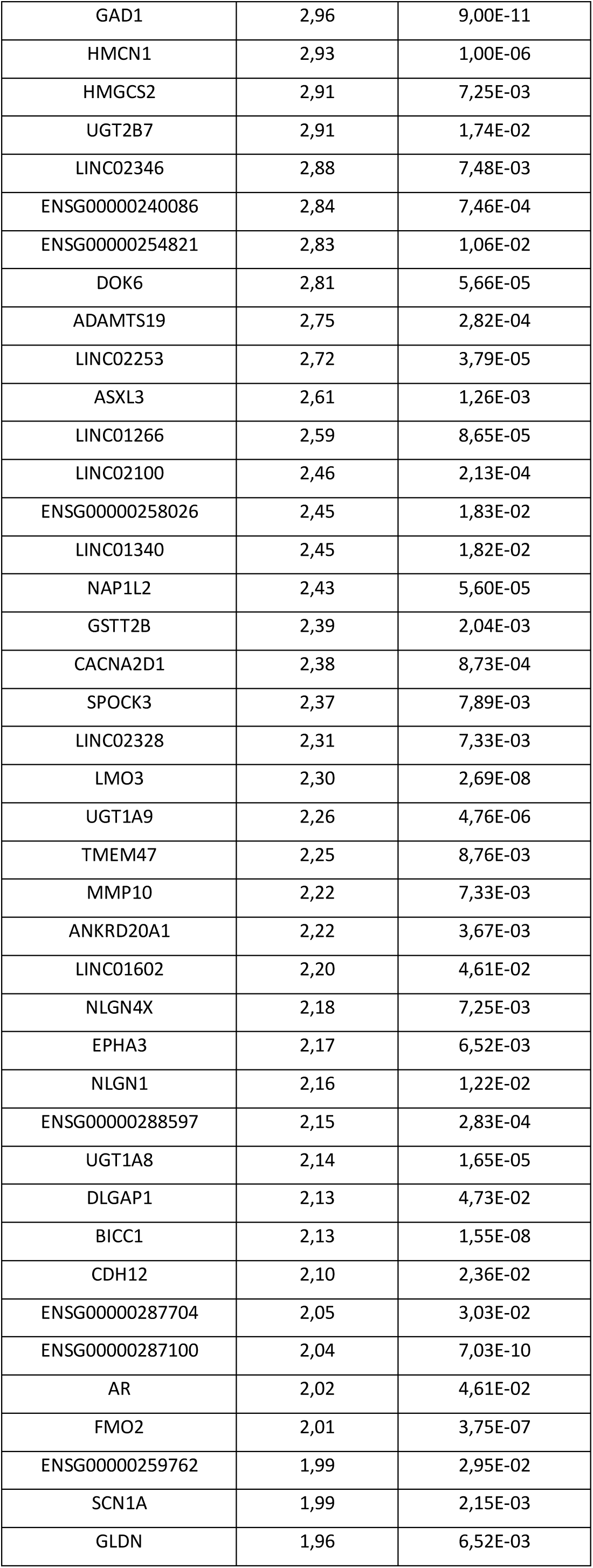
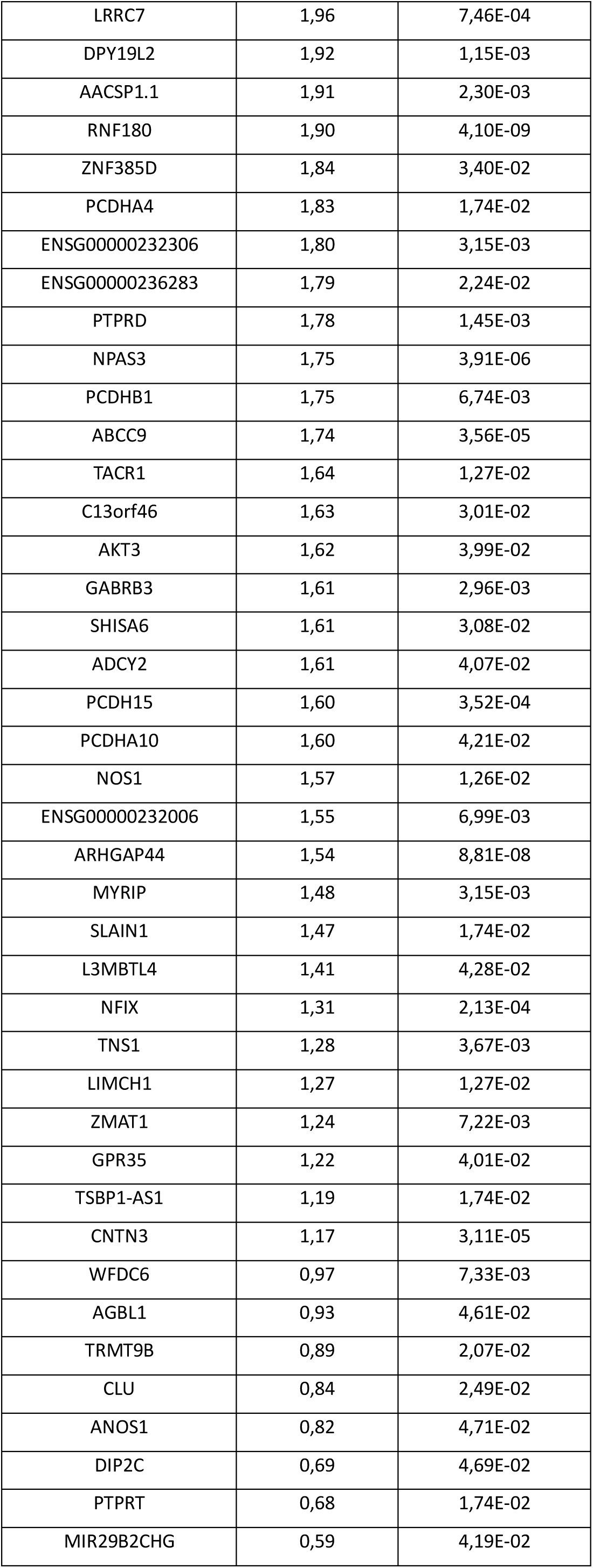
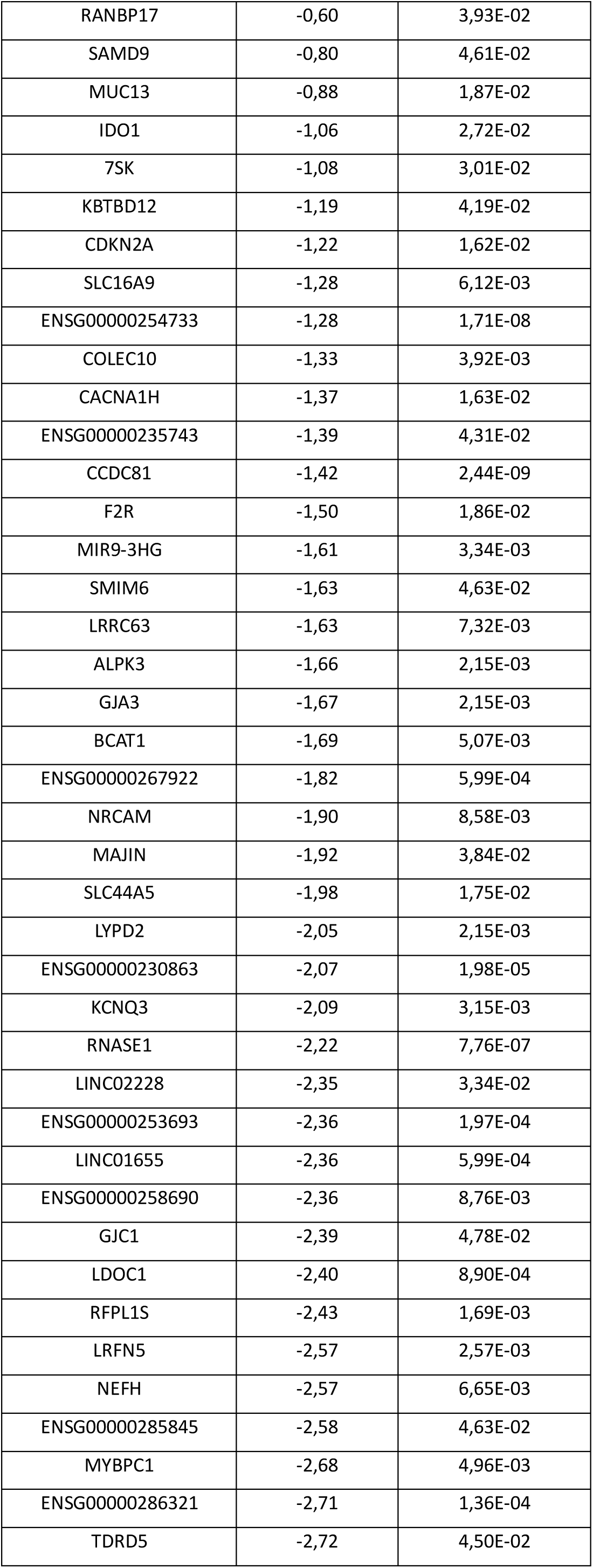
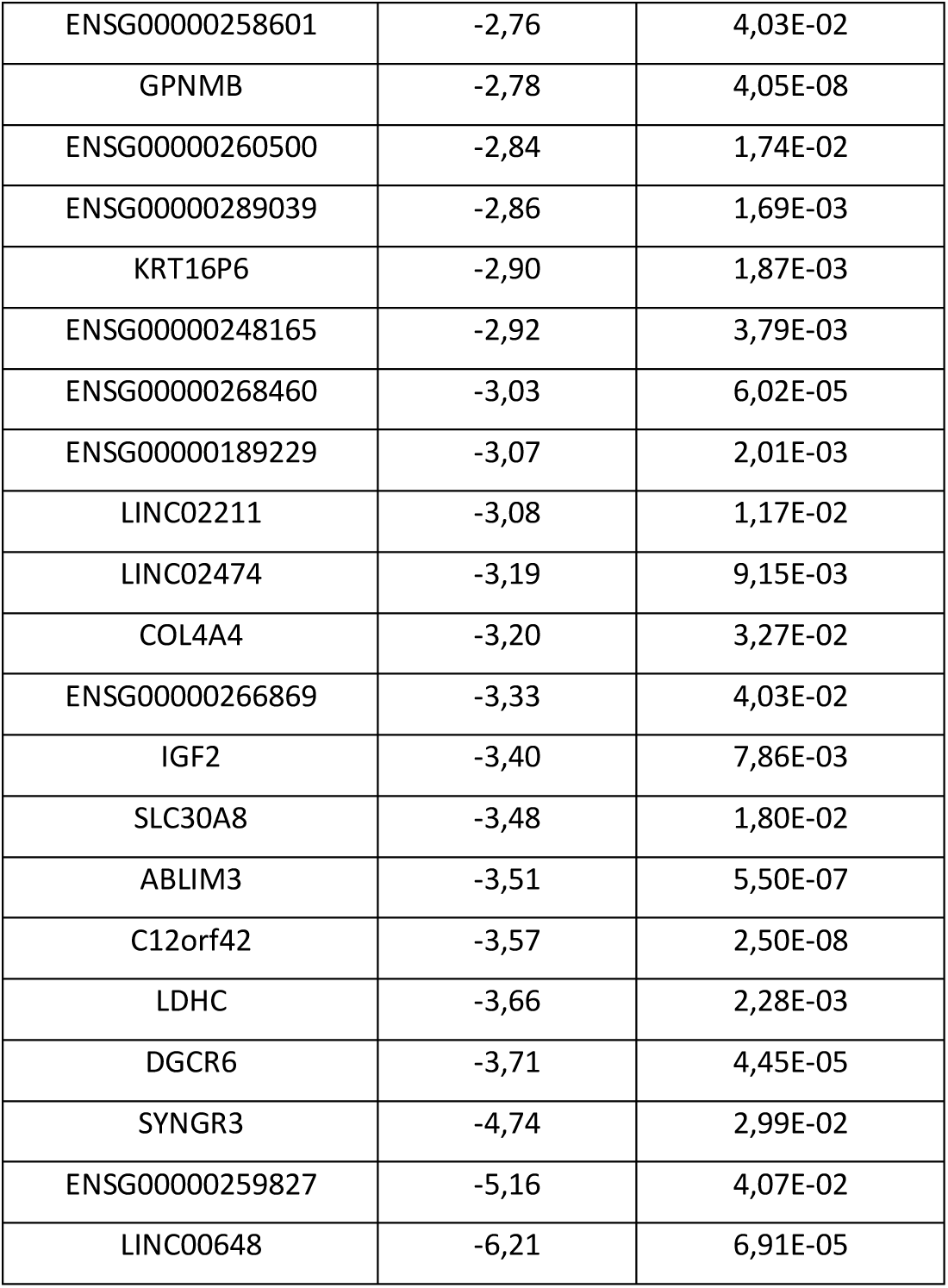
Pseudobulk differential gene expression analysis between adult and pediatric BE.

**Supplementary Table S2A.** Sample list barcoding for the Parse single cell experiments.

| sample_name | barcoding |
| --- | --- |
| 724_CT1 | A1-A2 |
| 724_RVA | A3-A4 |
| 724_RVC | A5-A6 |
| 724_ADV | A7-A8 |
| 724_CT2 | A9-A10 |
| 724_SAR | A11-A12 |

**Supplementary Table S2B.** Sample list barcoding for the Parse single cell experiments.

| sample_name | barcoding |
| --- | --- |
| V14_CTL | A1-A2 |
| V14_RVA | A3-A4 |
| V14_RVC | A5-A6 |
| V14_ADV | A7-A8 |
| V14_SAR | A9-A10 |
| 736_CTL | A11-A12 |
| 736_RVA | B1-B2 |
| 736_RVC | B3-B4 |
| 736_ADV | B5-B6 |
| 736_SAR | B7-B8 |
| 739_CTL | B9-B10 |
| 739_RVA | B11-B12 |
| 739_RVC | C1-C2 |
| 739_ADV | C3-C4 |
| 739_SAR | C5-C6 |
| V16_CTL | C7-C8 |
| V16_RVA | C9-C10 |
| V16_RVC | C11-C12 |
| V16_ADV | D1-D2 |
| V16_SAR | D3-D4 |
| V18_CTL | D5-D6 |
| V18_RVA | D7-D8 |
| V18_RVC | D9-D10 |
| V18_ADV | D11 |
| V18_SAR | D12 |

**Supplementary Table S3.**
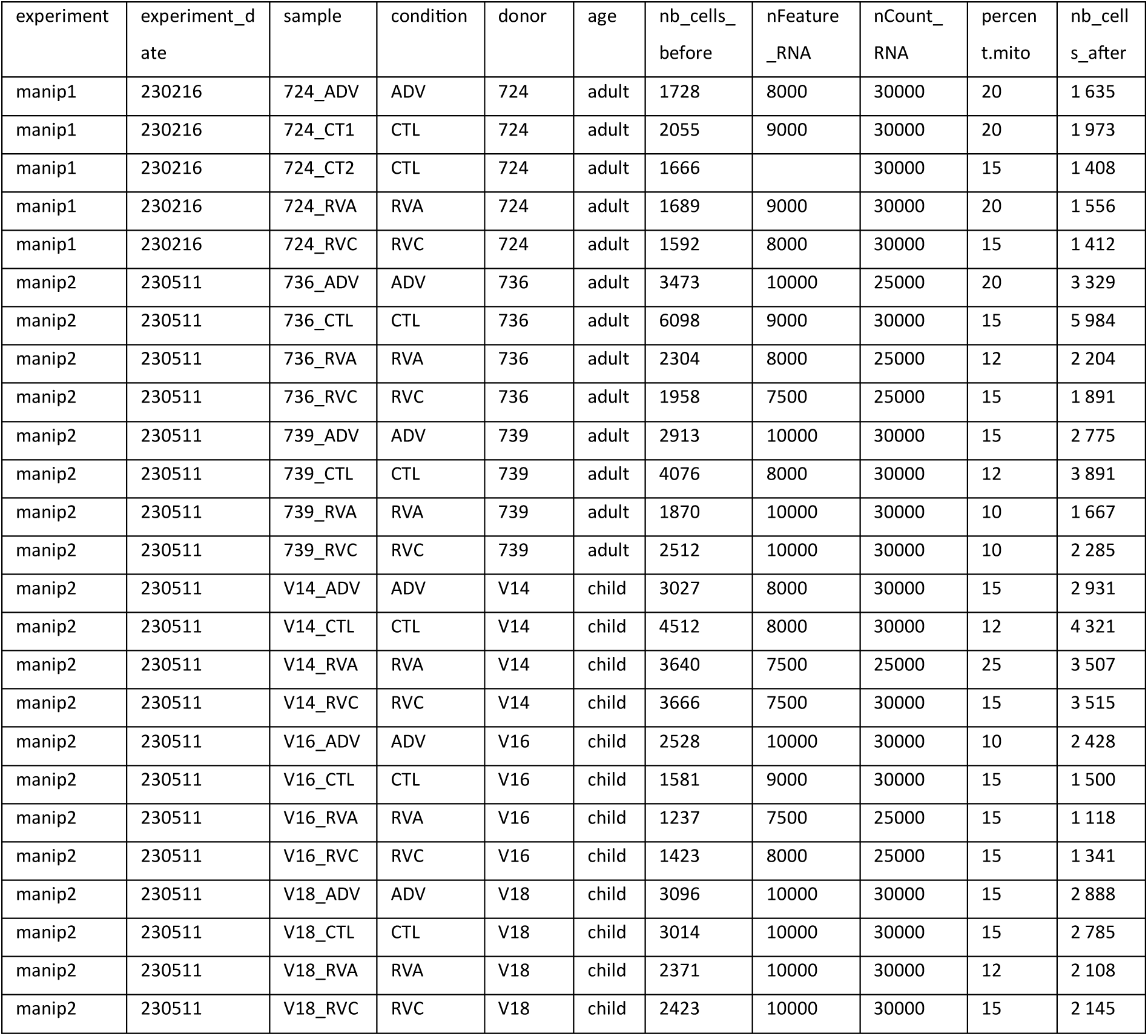
Metadata and threshold to filter outliers cells in the Parse single cell experiments.

**Supplementary Table S4.**
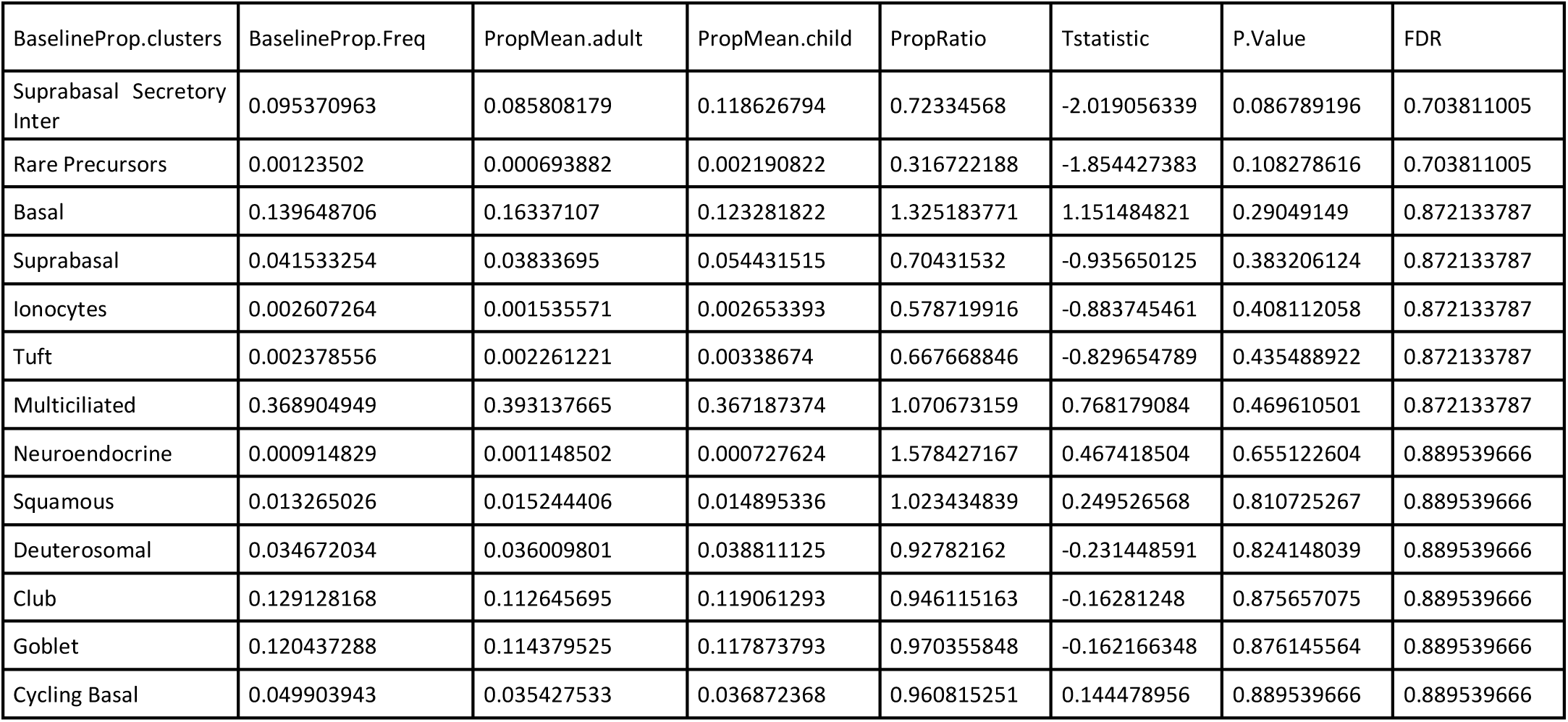
Proportion analysis using Propeller with asin transformation.

**Supplementary Table S5.** Metadata associated with the RNA-seq dataset.

| ID | Patient | Day_post_infection | condition | Cross_protection_GFP_Control | Cross_protection_neg_log10 |
| --- | --- | --- | --- | --- | --- |
| U798 | 867 | D1 | AdV | 25.84601859 | -1.412393652 |
| U795 | 867 | D1 | C | 1 | 0 |
| U796 | 867 | D1 | RVA | 0.131374027 | 0.881490487 |
| U797 | 867 | D1 | RVC | 0.070082894 | 1.154387973 |
| U820 | 867 | D5 | AdV | 0.109282122 | 0.961450881 |
| U818 | 867 | D5 | RVA | 0.052278616 | 1.281675918 |
| U819 | 867 | D5 | RVC | 0.010973937 | 1.959637537 |
| U834 | 867 | D12 | RVC | 1.049560853 | -0.021007624 |
| U832 | 867 | D12 | C | 0.611250523 | 0.213780756 |
| U833 | 867 | D12 | RVA | 0.175031368 | 0.756884113 |
| U850 | 868 | D1 | AdV | 7.598119266 | -0.880706106 |
| U847 | 868 | D1 | C | 1 | 0 |
| U848 | 868 | D1 | RVA | 0.000001 | 6 |
| U849 | 868 | D1 | RVC | 0.001536697 | 2.813411757 |
| U872 | 868 | D5 | AdV | 0.015375321 | 1.813175808 |
| U870 | 868 | D5 | RVA | 0.000629648 | 3.200902172 |
| U871 | 868 | D5 | RVC | 0.000561578 | 3.250589914 |
| U887 | 868 | D12 | AdV | 0.015574497 | 1.807585971 |
| U884 | 868 | D12 | C | 1 | 0 |
| U885 | 868 | D12 | RVA | 0.002674409 | 2.572772175 |
| V049 | 870 | D1 | AdV | 1.646063536 | -0.216446594 |
| V046 | 870 | D1 | C | 1 | 0 |
| V047 | 870 | D1 | RVA | 0.312154696 | 0.505630127 |
| V048 | 870 | D1 | RVC | 0.033667127 | 1.472793943 |
| V071 | 870 | D5 | AdV | 0.123786097 | 0.90732813 |
| V069 | 870 | D5 | RVA | 7.21131E-05 | 4.141985835 |
| V070 | 870 | D5 | RVC | 0.000697093 | 3.156709278 |
| AN744 | 902 | D1 | C | 1 | 0 |
| AN745 | 902 | D1 | RVA | 0.04005334 | 1.397361263 |
| AN746 | 902 | D1 | RVC | 0.01082325 | 1.96564231 |
| AN763 | 902 | D5 | AdV | 0.00523014 | 2.281486686 |
| AN761 | 902 | D5 | RVA | 0.00013505 | 3.869505411 |
| AN762 | 902 | D5 | RVC | 0.00179249 | 2.746543259 |
| AN779 | 902 | D12 | AdV | 0.00128173 | 2.892203451 |
| AN777 | 902 | D12 | RVA | 0.40090134 | 0.396962492 |
| AN778 | 902 | D12 | RVC | 0.03377453 | 1.471410686 |
| AN803 | 910 | D1 | AdV | 10.2934159 | -1.012559521 |
| AN800 | 910 | D1 | C | 1 | 0 |
| AN801 | 910 | D1 | RVA | 0.28066661 | 0.551809251 |
| AN802 | 910 | D1 | RVC | 1.16601402 | -0.066703772 |
| AN847 | 910 | D5 | AdV | 0.02806122 | 1.551893451 |
| AN844 | 910 | D5 | C | 1 | 0 |
| AN845 | 910 | D5 | RVA | 0.0522162 | 1.282194737 |
| AN846 | 910 | D5 | RVC | 0.11447704 | 0.941281609 |
| AN911 | 910 | D12 | AdV | 1.62380038 | -0.210532639 |
| AN908 | 910 | D12 | C | 1 | 0 |
| AN909 | 910 | D12 | RVA | 0.02436144 | 1.613297044 |
| AN910 | 910 | D12 | RVC | 0.02997195 | 1.523285001 |

